# SntB triggers the antioxidant pathways to regulate development and aflatoxin biosynthesis in *Aspergillus flavus*

**DOI:** 10.1101/2023.12.19.572357

**Authors:** Dandan Wu, Chi Yang, Yanfang Yao, Dongmei Ma, Hong Lin, Ling Hao, Wenwen Xin, Kangfu Ye, Minghui Sun, Yule Hu, Yanling Yang, Zhenhong Zhuang

## Abstract

The epigenetic reader SntB was identified as an important transcriptional regulator of growth, development, and secondary metabolite synthesis in *Aspergillus flavus*. However, the underlying molecular mechanism is still unclear. In this study, *sntB* gene deletion (Δ*sntB*), complementary (Com-*sntB*), and HA tag fused to *sntB* (*sntB*-HA) strains were constructed by using the homologous recombination method, respectively. Our results revealed that deletion of *sntB* inhibited the processes of mycelia growth, conidial production, sclerotia formation, aflatoxin synthesis, and ability to colonize host, and the defective phenotype of knockout strain Δ*sntB* could be restored in its complementary strain Com-*sntB*. Chromatin immunoprecipitation sequencing (ChIP-seq) of *sntB-*HA and WT, and RNA sequencing (RNA-seq) of Δ*sntB* and WT strains revealed that SntB played key roles in oxidative stress response of *A. flavus*. The function of *catC* gene (encode a catalase) was further analyzed based on the integration results of ChIP-seq and RNA-seq. In Δ*sntB* strain, the relative expression level of *catC* was significantly higher than in WT strain, while a secretory lipase encoding gene (G4B84_008359) was down-regulated. Under the stress of oxidant menadione sodium bisulfite (MSB), the deletion of *sntB* obvious down-regulated the expression level of *catC*. After deletion of *catC* gene, the mycelia growth, conidial production, and sclerotia formation were inhibited, while ROS level and aflatoxin production were increased compared to the WT strain. Results also showed that the inhibition rate of MSB to Δ*catC* strain was significantly lower than that of WT group and AFB1 yield of the Δ*catC* strain was significantly decreased than that of WT strain under the stress of MSB. Our study revealed the potential machinery that SntB regulated fungal morphogenesis, mycotoxin anabolism, and fungal virulence through the axle of from SntB to fungal virulence and mycotoxin bio-synthesis, i.e. H3K36me3 modification-SntB-Peroxisomes-Lipid hydrolysis-fungal virulence and mycotoxin bio-synthesis. The results of this study shed light into the SntB mediated transcript regulation pathways of fungal mycotoxin anabolism and virulence, which provided potential strategy for control the contamination of *A. flavus* and its aflatoxins.

## 1. Introduction

*Aspergillus flavus* is one of the common asexual species, a saprophytic fungus and the second largest pathogenic fungus after *Aspergillus fumigatus*, widely distributed in soil, air, water, plants, and agricultural products in nature [1]. Aflatoxins produced by *A. flavus* have strong toxicity, and is extremely harmful to human society. Animal and human health can be negatively affected by aflatoxins, which are carcinogenic, teratogenic, and mutagenic [2]. Among aflatoxins, AFB1 is the most frequently occurring and the most toxic and carcinogenic, which is converted to AFB1-8 and 9-epoxide in the liver and formed adducts with the guanine base of DNA and thus results in acute and chronic diseases in both human and household animals [3]. According to the Food and Agriculture Organization of the United Nations, 25% of the food crops in the world are contaminated with aflatoxins [4], which are often detected in grains, nuts, and spices [5, 6]. It is urgent to control the contamination of *A. flavus* and its main mycotoxin, AFB1.

In recent decades, the biosynthetic pathway of aflatoxins was investigated in detail benefit from the sequence of *A. flavus* genome [7]. This pathway consists of a complex set of enzymatic reactions [8–11]. In general, these enzymes are encoded by clusters of genes, which are regulated by cluster-specific genes: *aflR* and *aflS* [12, 13]. The initial stage of aflatoxins biosynthesis is catalyzed by polyketide synthase (PKSA) to form the polyketone backbone [14]. The synthesis of aflatoxins is additionally influenced by environmental stimuli such as pH, light exposure, nutrient availability, and the response to oxidative stress, potentially leading to the alteration of gene expression related to toxin biosynthesis [15–17]. Besides the biosynthetic pathway and its internal gene regulation, protein post-translational modifications (PTMs), an important mean of epigenetics, represent an important role in the regulation of aflatoxins synthesis, including 2-hydroxyisobutyrylation, succinylation, acetylation, and methylation [18–24], in which Snt2 (also called *sntB*, an epigenetic reader) is deeply involved. Despite advancements in the field, our understanding of the molecular mechanisms of aflatoxin production in *A. flavus* is still fragmentary.

The epigenetic reader encoded by *sntB* in *A. nidulans* was identified as a transcriptional regulator of the sterigmatocystin biosynthetic gene cluster and deletion of *sntB* gene in *A. flavus* results in loss of aflatoxin production [25], increasing global levels of H3K9K14 acetylation and impairing several developmental processes [20]. The homolog gene in yeast, SNTB coordinates the transcriptional response to hydrogen peroxide stress [26, 27]. In *Penicillium expansum*, SntB regulated the development, patulin and citrinin production, and virulence on apples [28]. In *A. nidulans*, SntB combined with a H3K4 histone demethylase KdmB, a cohesin acetyltransferase (EcoA), and a histone deacetylase (RpdA) to form a chromatin binding complex (KERS) and bound to regulatory genes and coordinated fungal development with mycotoxin synthesis [29]. In *A. flavus*, the KERS complex also consists of KdmB, RpdA, EcoA, and SntB and plays a key role in the fungal development and secondary metabolites metabolism [30]. *sntB* also regulated the virulence in *Fusarium oxysporum*, and respiration in *F. oxysporum* and *Neurospora crassa* [31, 32].

However, the specific regulatory mechanism of *sntB* in *A. flavus* remains unclear. In this study, we identified the regulatory network of *sntB* by Chromatin immunoprecipitation sequencing (ChIP-seq) and RNA sequencing (RNA-seq), which shed light on its impact on fungal biology.

## Experimental Procedures

### *A. flavus* strains, media, and culture conditions

*A. flavus* Δ*ku70* Δ*pyrG* was used as a host strain, for genetic manipulations. All strains used in this study are listed in the **Table 1**. potato dextrose agar (PDA, 39 g/L, BD, Difco, Franklin, NJ, USA), complete medium (CM, 6 g/L tryptone, 6 g/L yeast extract, 10 g/L glucose), and potato dextrose (PDB, 24 g/L, BD, Difco, Franklin, NJ, USA) were used for mycelial growth and sporulation determination, sclerotia production, and mycotoxin production analysis, respectively. All experiments were technically repeated three times and biologically repeated three times.

**Table 1.** *A. flavus* strains used in this study.

| Strain name | Related genotype | Source |
| --- | --- | --- |
| *A. flavus* CA14 | $\Delta ku70, \Delta pyrG$ | Kindly presented from Prof.<br>Chang<sup>[33]</sup> |
| Wild-type (WT) | $\Delta ku70, \Delta pyrG::AfpyrG$ | This study |
| $\Delta sntB$ | $\Delta ku70, \Delta sntB::AfpyrG$ | This study |
| Com-*sntB* | $\Delta ku70, \Delta pyrG; \Delta sntB::AfpyrG::$<br>*sntB* | This study |
| *sntB*-HA | $\Delta ku70$, *sntB*-HA::*Afp<sub>pyrG</sub>* | This study |
| $\Delta catC$ | $\Delta ku70$, $\Delta catC$::*Afp<sub>pyrG</sub>* | This study |

### The construction of mutant strains

All mutant strains, including *sntB* and *catC* gene knock-out strain (Δ*sntB* and Δ*catC*), the complementation strain for the Δ*sntB* strain (Com-*sntB*), and HA tag fused to *sntB* strain (*sntB*-HA), were constructed following the protocol of homologous recombination [34] and the detail protocol was as described in our previous study [35]. The related primers were listed in Table 2. The constructed strains were confirmed by diagnostic PCRs and by southern blot [36]. The construction of *sntB*-HA was further determined by western blot with anti-HA antibody as descripted previously [35].

**Table 2.**
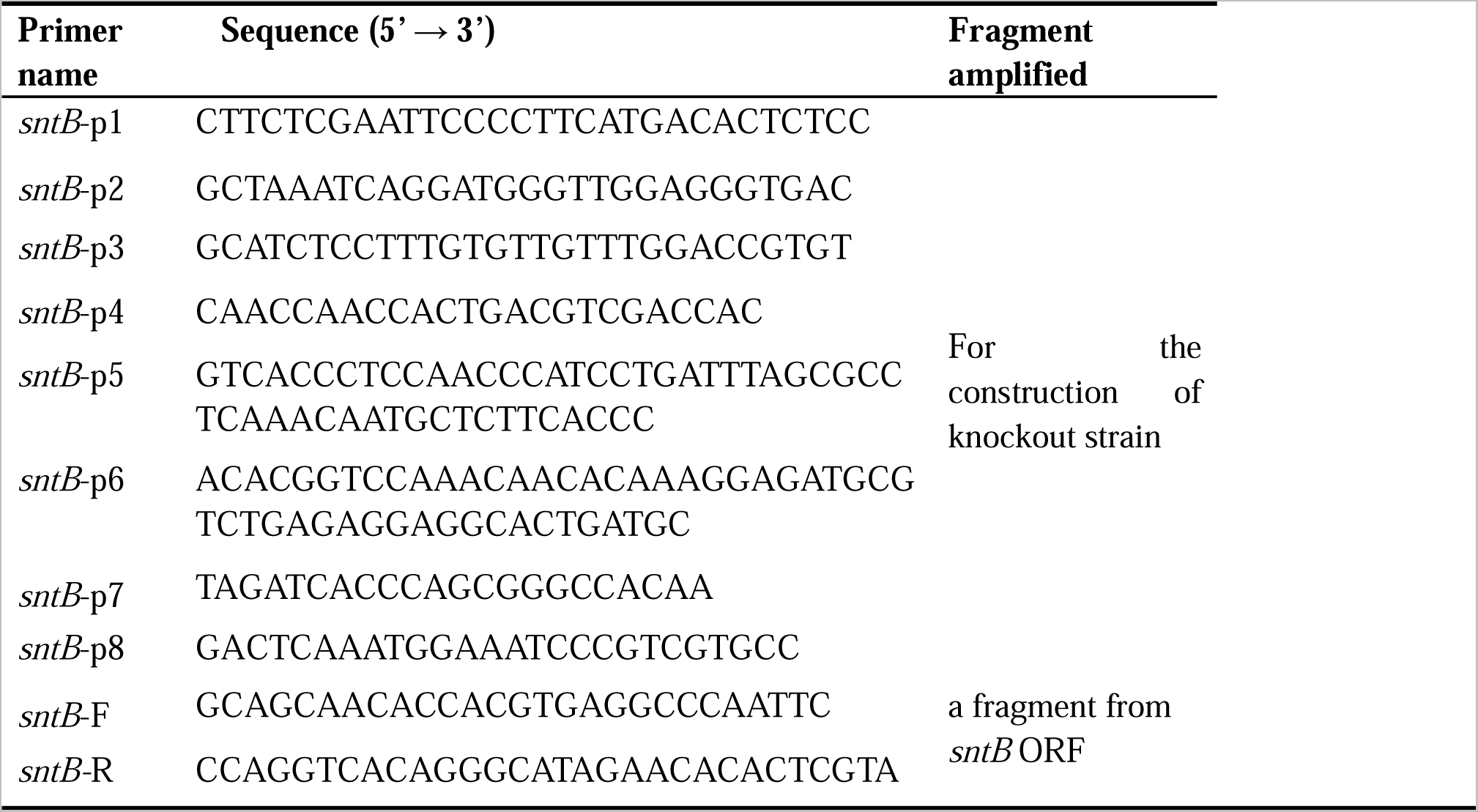
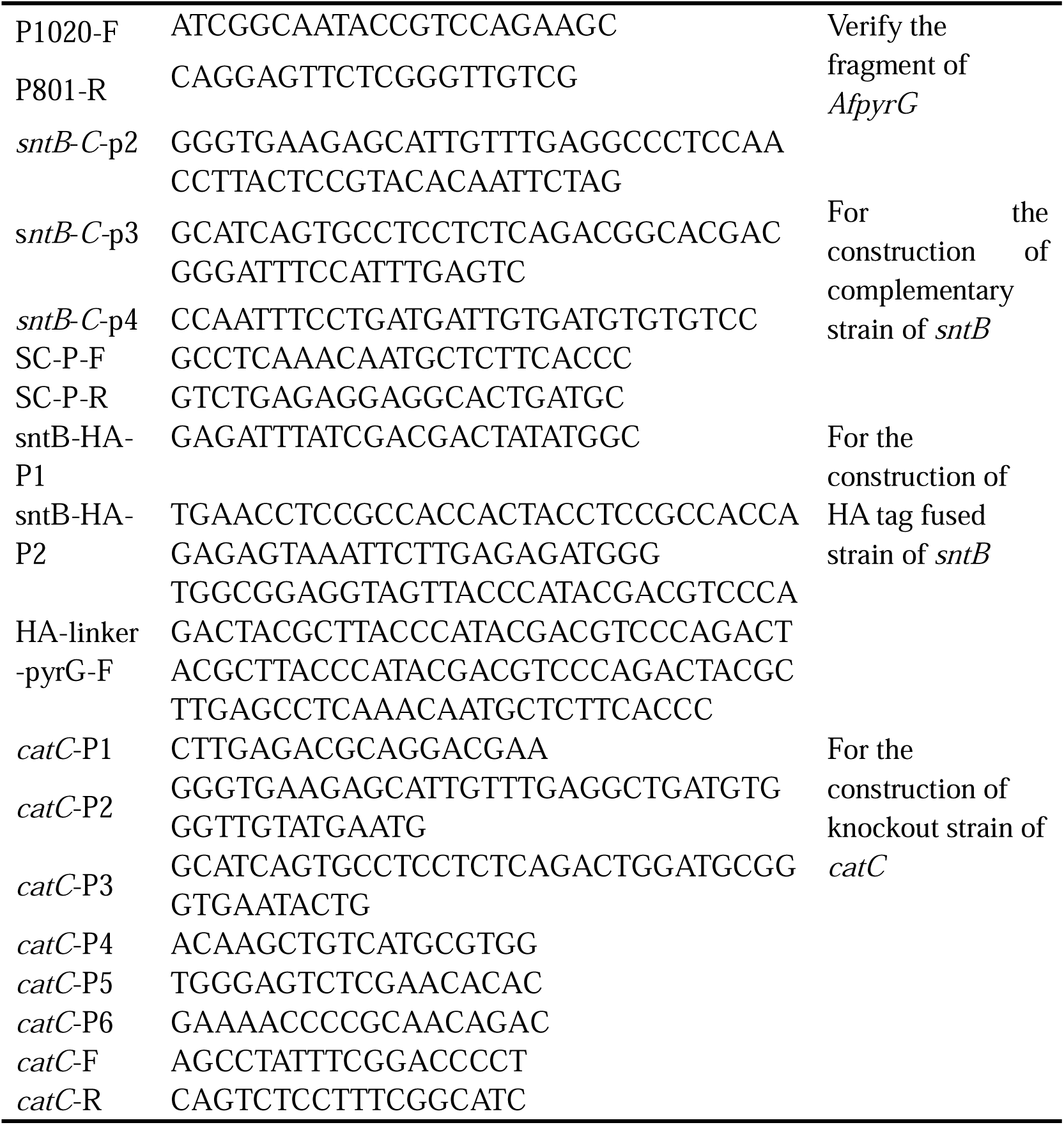
Primers used for strain construction in this study.

### Phenotypic analysis and aflatoxin analysis

The spores (10^7^ conidia/mL) of wild type (WT), Δ*sntB*, and Com-*sntB* strains were used. The details of experiment were according to our previous study [37]. Hyphal septum was stained according to the descripted method [38]. Each fungal strain was evaluated on four plates, and each experiment was repeated three times.

### Fungal colonization on crop kernels

According to our previous experimental protocol [37], the colonizing ability of WT, Δ*sntB*, and Com-*sntB* fungal strains on peanut and corn kernels was analyzed. The crop kernels were disinfected with 0.05% sodium hypochlorite and soaked for 30 min in a solution containing 10^5^ conidia/mL fungal spores. Afterwards, the seeds were placed in a petri dish and cultured at 29°C for 6 d. Finally, the number of conidia was calculated and AFB1 product was analyzed by TLC.

### Animal invasion assay

The animal invasion assay using silkworms was conducted according to a previous study [34, 39]. Silkworms (*Bombyx mori*) were randomly separated into four groups (10 larvae/group) when silkworm larva reach about 1 g in weight. Each silkworm was injected with 5 µL saline, or 5 µL conidial suspension (10^6^ spores/mL) from WT, Δ*sntB*, and Com-*sntB* strains. The survival rate of silkworms was calculated. Dead silkworms were transferred into fresh 9 cm Petri dishes and cultivated for 5 d in the dark. The conidia number and AFB1 production from each group were measured.

### RNA-seq analysis

To reveal the potential complex regulatory network of the *sntB*, RNA-seq analysis was carried out on the WT and Δ*sntB* strains by APPLIED PROTEIN TECHNOLOGY, Shanghai (www.aptbiotech.com) [40]. Data processing was according to a previous study [41]. Differentially expressed gene (DEGs) were assigned as genes with |log2FoldChange|>1 and adjusted *P*-adj<0.05. Gene Ontology (GO) and Kyoto Encyclopedia of Genes and Genomes (KEGG) pathways were used to analyze the functions of DEGs.

### ChIP-seq and data analysis

ChIP-seq analysis was carried out on the WT and *sntB*-HA strains. The conidia (10^4^/ml) for each strain were inoculated in 100 mL PDB shaking at 180 rpm under 29°C for 72 h, and subjected to ChIP-seq analysis by Wuhan IGENEBOOK Biotechnology Co., Ltd (http://www.igenebook.com). ChIP experiment was according to a previous study [34]. Raw sequencing with low-quality reads were discarded, and reads contaminated with adaptor sequences trimmed were filtered by Trimmomatic (v0.36) [42]. The clean reads were mapped to the reference genome of *A. flavus* by Burrows-Wheeler Alignment tool (BWA, version 0.7.15) [43]. MACS2 (v2.1.1) and Bedtools (v2.25.0) were used for peak calling and peak annotation, respectively. Differential binding peaks were identified by Fisher’s test with q value <0.05. HOMER (version3) was used to predict motif occurrence within peaks with default settings for a maximum motif length of 12 base pairs [44]. Genes less than 2000 bp away were associated to the corresponding peak. GO and KEGG enrichment analyses of annotated genes were implemented in EasyGO [45] and KOBAS (v2.1.1) [46], with a corrected *P*-value cutoff of 0.05.

### Quantitative RT-PCR (qRT-PCR) analysis

The fungal spores (10^6^/mL) were cultured in PDB medium for 48 h, and then mycelium was ground into powder with liquid nitrogen. Total RNA was prepared by TRIpure total RNA Extraction Reagent (Bestek, China) according to the protocol used by Zhang [47]. qRT-PCR was performed according to previously described [48], and the primers were shown in **Table 3**.

**Table 3.**
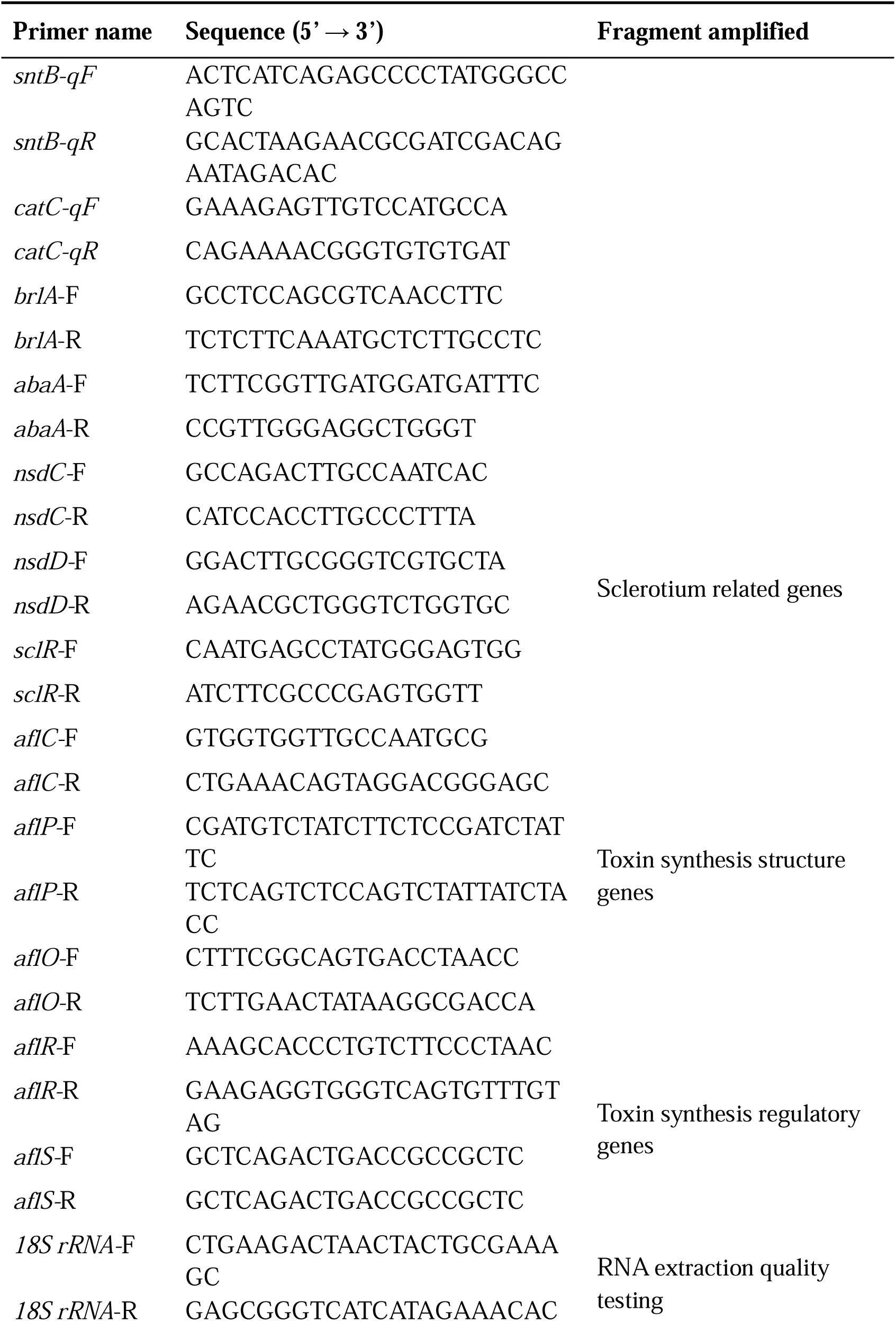
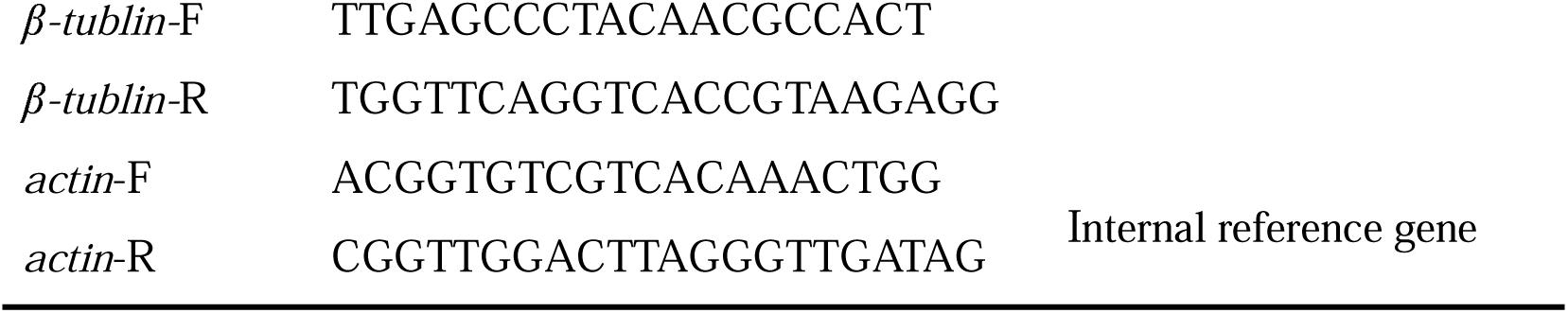
Primers used for RT-qPCR in this study.

### Oxidative stress assays

To evaluate the role of *sntB* in fungal resistance to oxidative stress, a series concentration (0, 0.12, 0.24, and 0.36 mM) of menadione sodium bisulfite (MSB) were added to the medium. 10^6^ fungal spores for each strain were inoculated on the medium and cultured in dark at 37L. The diameters of colonies were measured 3 d after inoculation and the inhibition rate was calculated as previously descripted [35]. The AFB1 product was analyzed by TLC after the strains were cultured in YES medium in dark at 29L for 7 d.

### Reactive oxygen species assay

With the instructions provided in the user’s manual, the intracellular ROS production was measured using a ROS assay kit (S0033S, Beyotime Institute of Biotechnology, China). After harvest, the mycelia were incubated with 10 µM 6-carboxy-2’,7’-dichlorodihydrofluorescein diacetate (DCFH-DA) and 50 g/mL Rosup for 30 min. With SpectraMax Imaging Cytometer (Molecular Devices, Sunnyvale, Calif.) at emission wavelength of 525 nm and excitation wavelength of 488 nm, fluorescence signals of intracellular ROS production were acquired.

### Statistical analysis

All data in this study were expressed as mean ± standard deviation. The statistical analysis was performed using the software GraphPad Prism8 (GraphPad Software, La Jolla, CA, USA). The difference was considered to be statistically significant when *P* < 0.05.

## Results

### The phenotype of SntB in *A. flavus*

The role of SntB in *A. flavus* has been previously characterized by analyzing both Δ*sntB* and overexpression *sntB* genetic mutants [20]. To further investigate the intrinsic mechanism of this regulator on the development and aflatoxin biosynthesis in *A. flavus*, the *sntB* deletion strain (Δ*sntB*) and the complementary strain (Com-*sntB*) were constructed by the method of homologous recombination and verified by diagnostic PCR and southern blotting (Figure S1). The expression levels of *sntB* in WT, Δ*sntB*, and Com-*sntB* strains was further detected by qRT-PCR and the result showed that the expression of *sntB* was absent in the gene-deletion strain, and it fully recovered in the Com-*sntB* strain (Figure S1), which reflected that the Δ*sntB* and Com-*sntB* strains had been successfully constructed, and could be used in the subsequent experiments of this study.

The phenotype analysis of this study revealed that the deletion of *sntB* gene significantly inhibited the growth of mycelium, hyphae morphology, the length of fungal cell (between two adjacent septa), the number of conidiation, sclerotium formation, and aflatoxin biosynthesis, while the above phenotypes of both development and mycotoxin bio-synthesis were fully recovered in the Com-*sntB* strain (Figure 1). To reveal the signaling pathways of *sntB* in conidiation, sclerotium formation, and aflatoxin biosynthesis, qRT-PCR analysis was performed to assess the expression levels of sporulation related transcriptional factor genes, *steA*, *wetA*, *fluG*, and *veA,* sclerotia formation related transcriptional factor genes, *nsdC*, *nsdD*, and *sclR* [49, 50], and the AFs synthesis gene cluster structural genes *aflC*, *aflR,* and *aflP*, and the main regulatory gene *aflR* and *aflS*. As shown in Figure S2, the relative expression levels of these genes were significantly lower in the Δ*sntB* strain compared to that of the WT strain, and recovered in the Com-*sntB* strain. These results indicated that SntB regulates the conidiation, sclerotium formation, and aflatoxin biosynthesis by the canonical signaling pathways mediated by these regulators.

**Figure 1.**
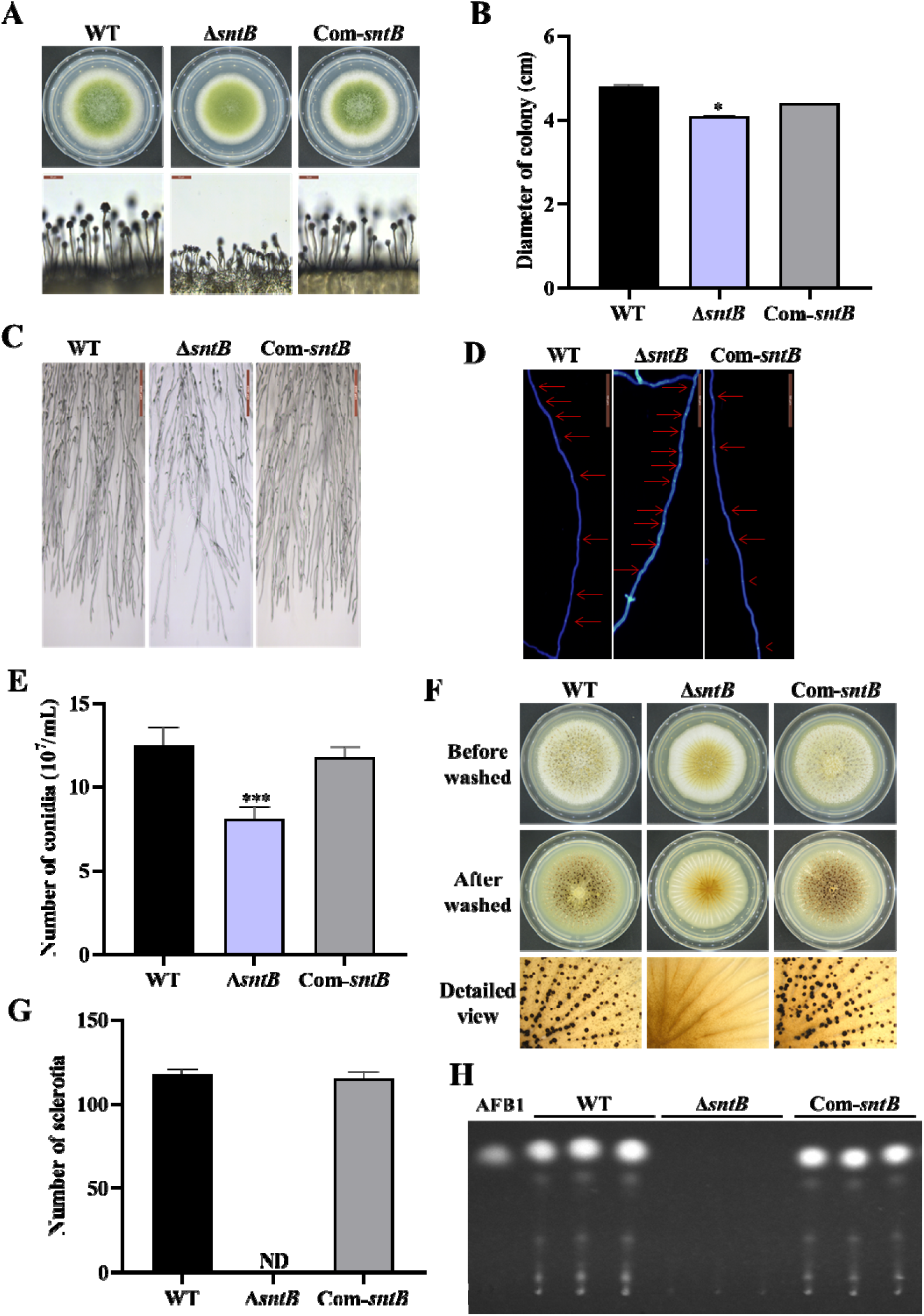
The functions of SntB in *A. flavus*. (A) The colonies of WT, Δ*sntB*, and Com*-sntB* strains grown on PDA at 37°C in dark for 4 d. (B) The colony diameter statistics of the above fungal strains. (C) Microscopic examination revealed the difference in mycelia of each fungi strain at 37L in dark, scale=200 μm. (D) Microscopic examination of the hyphal septum of each strain at 37L in dark, scale=50 μm. (E) The spore production statistics. (F) All the above fungal strains were point-inoculated on CM medium and grown for 7 d at 37L. (G) The number of sclerotia of the above fungal strains. ND=Not detectable. (H) AFB1 production of the above fungal strains was detected by TLC after the strains incubating at 29L in PDB medium for 7 d.

### SntB plays important roles in virulence of *A. flavus* to both plant and animal hosts

In order to explore the effect of SntB on the fungal colonization ability, peanut seeds and maize kernels were infected with spore solution of each fungal strain. Compared with WT, the conidiation yield of Δ*sntB* on the infected host was significantly reduced (*P*<0.001) and no AFB1 could be detected on the hosts infected by Δ*sntB*, while in the Com*-sntB* strain, the capacity to produce conidia and AFB1 on both crop kernels was recovered (Figure 2A-2C and Figure S3A). The role of *sntB* in fungal virulence to animals was also investigated. As Figure 2D-2E shown, the survival rate of silkworms injected by spores of Δ*sntB* strain was significantly higher than that from WT infected larvae. There was less fungal mycelium, conidia, and AFB1 production on the dead silkworms injected by Δ*sntB* compared to the silkworms from the WT injection groups, but when the gene was reintroduced (i.e. the Com-*sntB* group), similar to what found in the WT group, the survival rate of silkworms obviously dropped and more fungal mycelium, conidia, and AFB1 produced on the dead silkworms (Figure 2F-2H). All the above results revealed that SntB plays an essential role in virulence of *A. flavus*.

**Figure 2.**
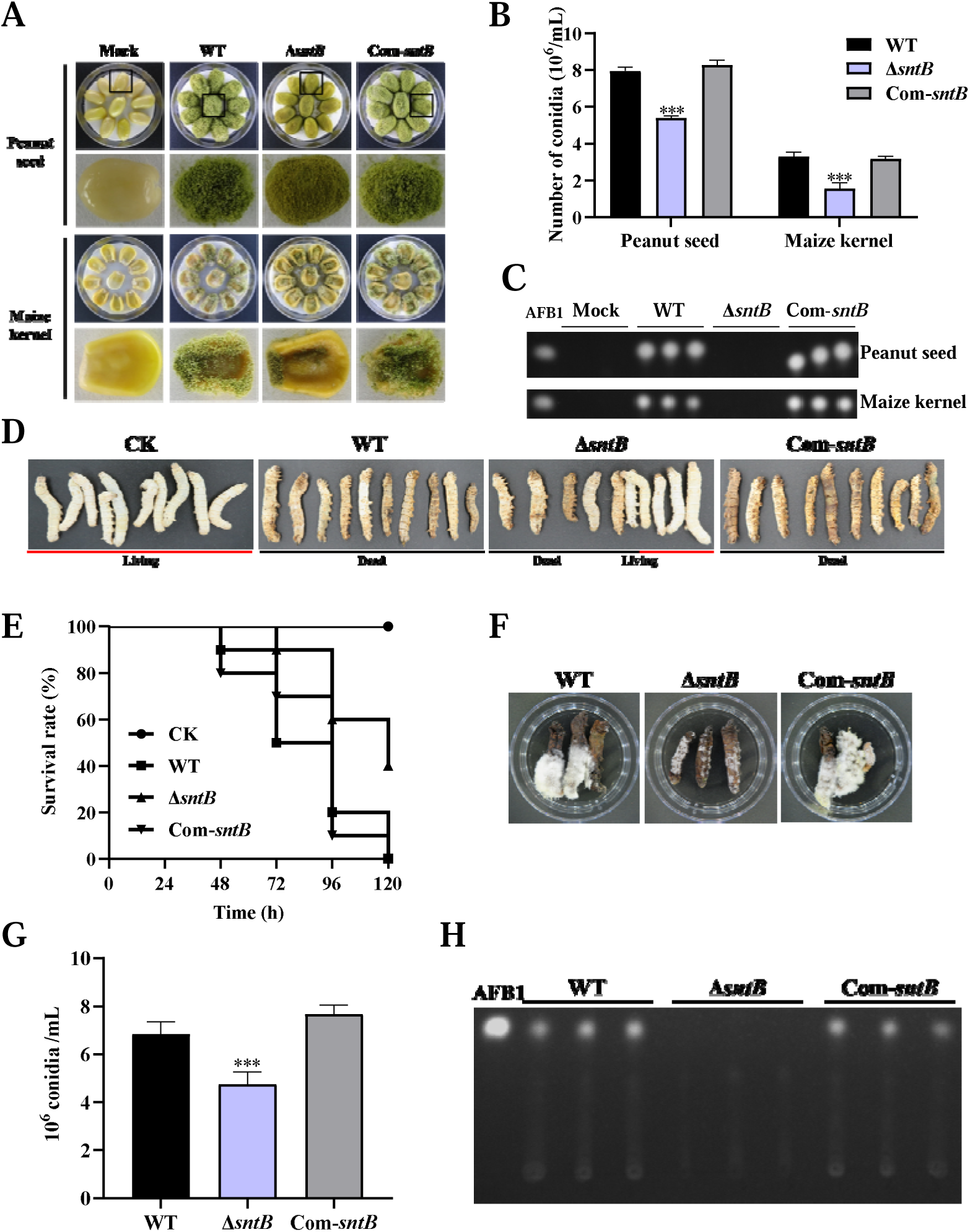
The role of SntB on the ability of *A. flavus* to colonize host. (A) Phenotypic of corn and peanut kernels colonized by Δ*sntB*, Com-*sntB*, and WT strains at 29°C in dark for 7 d. (B) Statistical of the number of conidia on the surface of peanut and maize kernels. (C) TLC analysis to detect the yield of AFB1 in kernels infected by the above fungal strains after 7 d incubation. (D) Photographs of the silkworms infected by the above fungal strains. (E) The survival rate of silkworms in 5 d after injection of the above strains. (F) Photographs of the dead silkworms infected by *A. flavus* after 6 d incubation. (G) The spore production statistics of the above fungal strains on the dead silkworms shown in (F). (H) TLC analysis of AFB1 levels produced in infected dead silkworms in (F).

The capacity of fungal infection is closely related to secreted hydrolases, such as amylase, lipase, protease, etc. In order to explore the effect of SntB on the activity of hydrolases, the activities of amylase in the above each fungal strain were further determined. The results showed that the colonies of Δ*sntB* produced almost no degradation transparent circle after adding iodine solution compared with that of WT and Com*-sntB*, which indicated that the activity of α-amylase in Δ*sntB* strain was significantly reduced (*P*<0.001) (Figure S3B and S3C). This suggests that SntB plays an important role in the fungal pathogenicity by changing the hydrolases activity of *A. flavus*.

### SntB chords global gene expression

To explore the downstream signaling pathways regulated by SntB, samples with three biological replicates of WT and Δ*sntB* strains were submitted for RNA-Seq. In the assay, the bases score Q30 was more than 93.19% (Table S1) and mapping ratio was from 95.17% to 95.80% (Table S2). To further confirm the quality of RNA-Seq, PCA (Principal Components Analysis) and pearson correlation analysis were performed. Correlation analysis revealed that the samples were clustered by groups (Figure 3A). A plot of PC1 (47.70%) and PC2 (23.20%) scores showed a clear separation between the groups (Figure 3B). A total of 1,446 and 1,034 genes were significantly up- and down-regulated, respectively, in the Δ*sntB* compared to the WT strain (Figure 3C and Supplementary Table S3). GO enrichment analysis identified 93 enriched GO terms (*P*<0.05) (Figure 3D and Supplementary Tables S4). In the biological process category, the most enriched terms were “oxidation-reduction process (GO: 0055114)”. In the molecular function category, “catalytic activity (GO: 0003824),” “oxidoreductase activity (GO: 0016491),” and “cofactor binding (GO: 0048037)” were the most significantly enriched terms. Whereas terms associated with “Set3 complex (GO: 0034967)”, “mitochondrial crista junction (GO: 0044284)” and “extracellular region (GO: 0005576)” were significantly enriched in the cellular component category. Additionally, all the DEGs were mapped according to the KEGG database, and 42 significantly enriched pathways were identified (*P*<0.05) (Supplementary Table S5). Among them, “metabolic pathways (ko01100),” “aflatoxin biosynthesis (ko00254),” and “microbial metabolism in diverse environments (ko01120)” were the most significantly enriched (Figure 3E and Supplementary Table S5).

**Figure 3.**
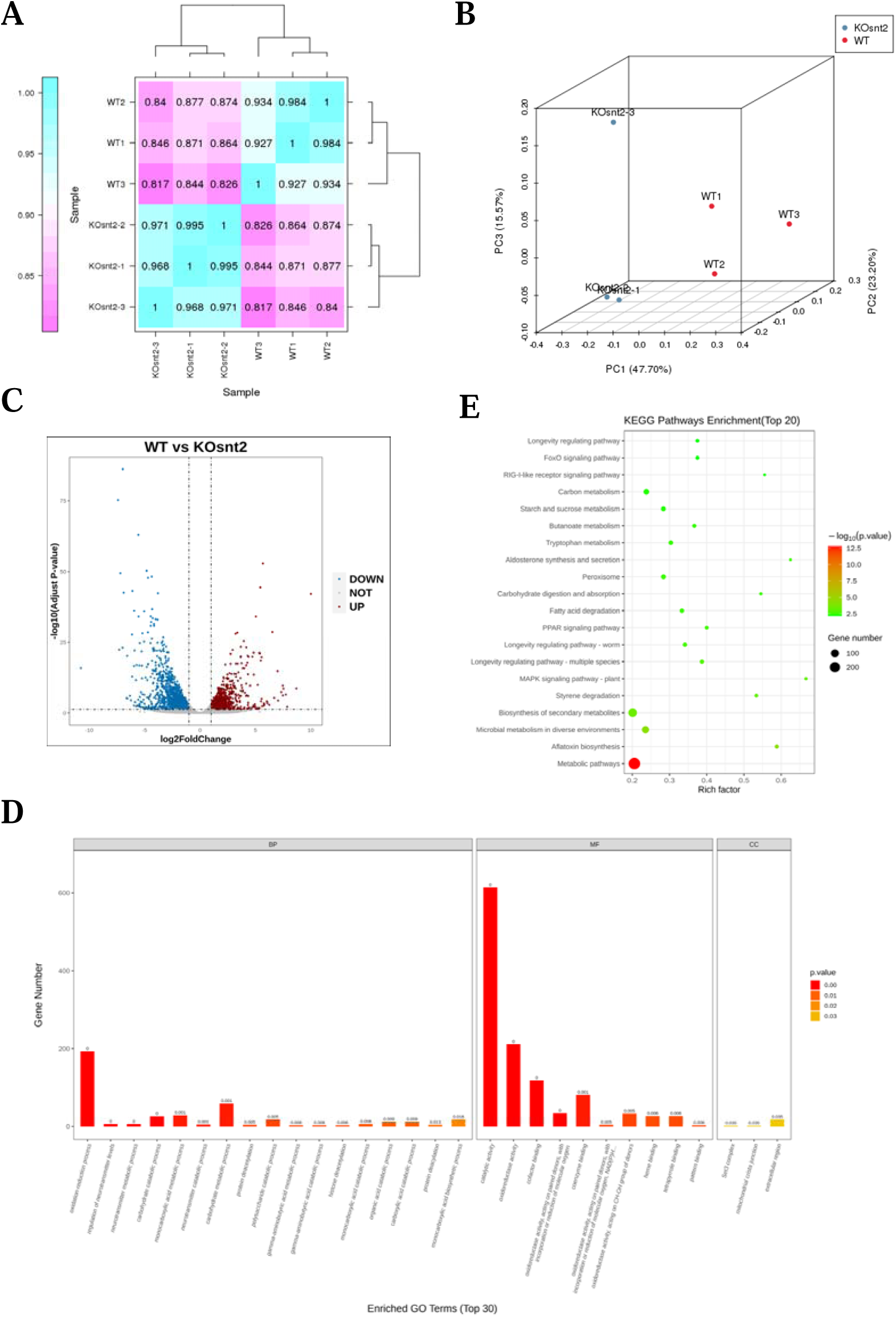
SntB chords global gene expression in *A. flavus*. (A) The pearson correlation results shown by heatmap. (B) Principal component analysis (PCA) on six fungal samples, including three Δ*sntB* (KOsnt2) and three WT samples. (C) Volcano map reflecting the distribution of the differentially expressed genes. (D) Gene Ontology (GO) analyses of the differentially expressed genes. (E) Kyoto encyclopedia of genes and genomes (KEGG) analyses of the differentially expressed genes.

### Characterization the binding regions of SntB by chromatin immunoprecipitation sequencing (ChIP-seq)

To characterize the chromatin regions targeted by SntB, ChIP-seq were carried out with both HA tag fused *sntB* strain (*sntB*-HA) and WT strain. The *sntB*-HA strain was constructed by homologous recombination through fused HA to the 3’ end of *sntB* (Figure 4A). In the ChIP-seq assay, more than 94.66% of bases score Q30 and above in each sample (Table S6), and reaching 52.50% to 94.48% of mapping ratio (Table S7). The PCA (Figure S4A) and heat map (Figure S4B) reflected that the quality of samples was competent for subsequent analysis. There were 1,510 up-enriched differently accumulated peaks (DAPs) in *sntB*-HA fungal strain compared to the WT strain, which were distributed on the whole *A. flavus* genome (Figure 4B, and Table S8). Most of the up-enriched peaks were located in the promoter (82.85%) region (Figure 4C). To determine binding regions of SntB, we used the HOMER known and *de novo* motif discovery algorithm. Motifs were sorted based on *p*-values and the top 5 enriched known motifs were shown in Figure 4D. The results consisted of motifs derived from previously-published ChIP-Seq experiments on Cbf1, bHLHE40, NFY, Usf2, and USF1 (Table S9). However, the most enriched *de novo* motif was NFYA (1e-97) (Figure 4E). The genes of the DAPs were further subjected to GO and KEGG analysis. The most strikingly enriched GO terms in the biological process category were “cell communication (GO:0007154)”, “response to stimulus (GO:0050896)”, and “response to external stimulus (GO:0009605)”. Whereas terms associated with “DNA-binding transcription factor activity (GO:0003700)”, “DNA-binding transcription factor activity, RNA polymerase II-specific (GO:0000981)”, and “sequence-specific DNA binding (GO:0043565)” were the most significantly enriched molecular function category (Figure 4F and Table S10). And these genes were mostly enriched in “Methane metabolism” and “MAPK signaling pathway - yeast” pathways (*P* value < 0.05) (Figure 4G and Table S11).

**Figure 4.**
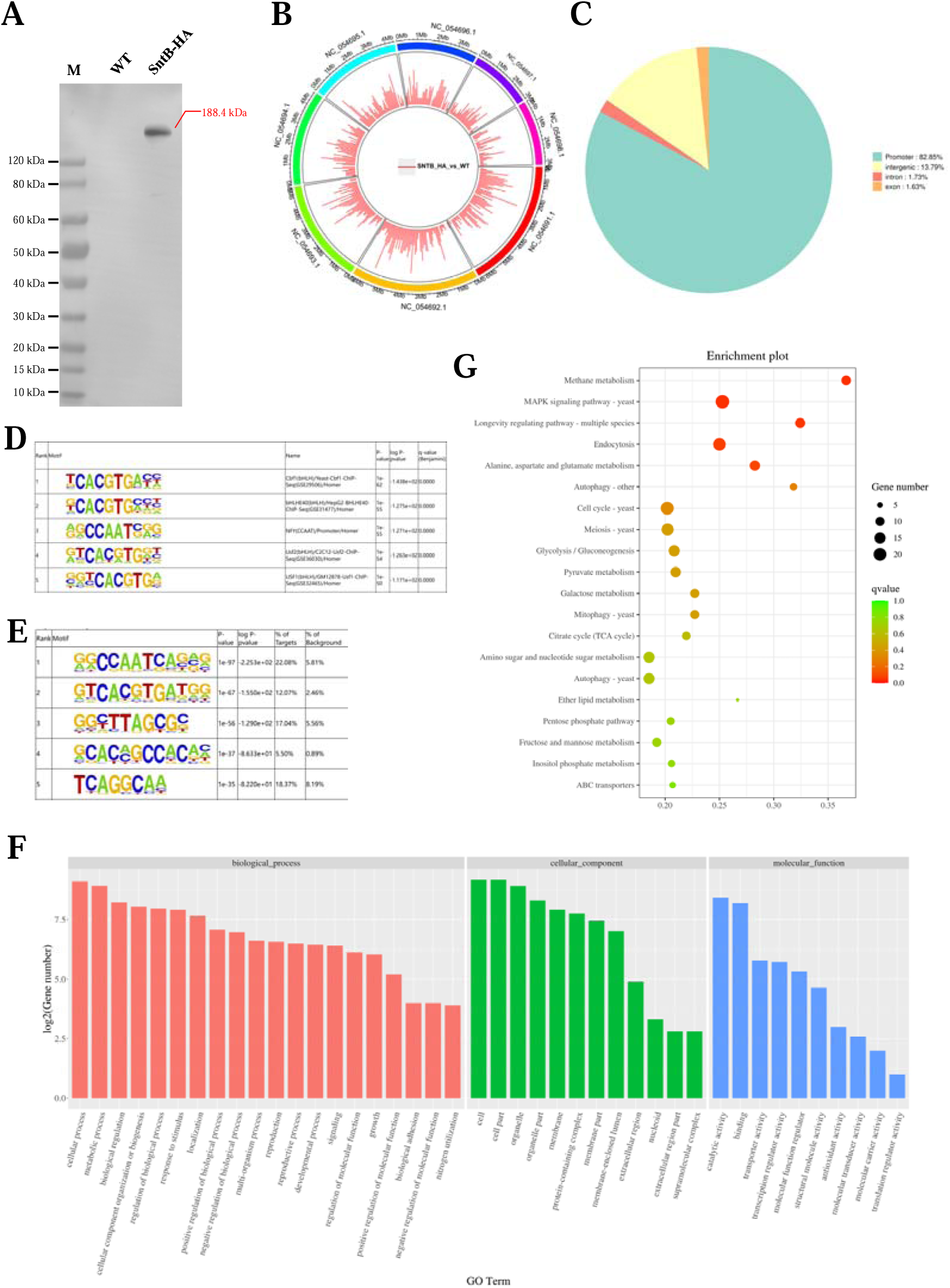
Characterization the binding regions of SntB. (A) Verification of the construction of *sntB*-HA strain using western blot. M means the protein marker of PAGE-MASTER Protein Standard Plus (GenScript USA, MM1397). (B) The distribution of differently accumulated peaks on the genome. (C) Vennpie map of the differently accumulated peaks distribution on gene functional elements. (D) Enrichment of known motifs showing the top-ranked motif logos. (E) Enrichment of *de novo* motifs showing the top-ranked motif logos. (F) GO analyses of the of the differently accumulated peaks related genes. (G) KEGG analyses of the differently accumulated peak related genes.

### Integration of the results of ChIP-seq and RNA-seq assays

After overlapping the results from both different sequence methods (ChIP-seq and RNA-seq), 238 DEGs were found (Figure 5A). According to the GO annotation, these DEGs were significantly enriched in 8 GO terms, including “cellular response to reactive oxygen species (GO:0034614)”, “reactive oxygen species metabolic process (GO:0072593)”, and “cellular response to oxygen-containing compound (GO:1901701)” (Figure 5B). It was further noted that the DEGs were significantly assigned to “carbon metabolism (afv01200)”, “peroxisome (afv04146)”, and “glyoxylate and dicarboxylate metabolism (afv00630)” KEGG pathways (Figure 5C and Table S12-S13). These results revealed that SNTB is essential for *A. flavus* to maintain the homeostasis of intracellular reactive oxygen species. Studies had shown that SNTB could response to oxidative stress in yeast [27] and *Magnaporthe oryzae* [51]. As Figure 5D and Figure 5E showed, due to the deletion of the *sntB* gene, Δ*sntB* exhibited a severe MSB sensitivity phenotype compared to that of the WT strain, and the phenotype recovered in the complementary strain (Com-Δ*sntB*). The results showed that the inhibition rate of oxidant MSB to Δ*sntB* would be significantly enhanced with the increase of MSB concentration. These results showed that SntB deeply participates in the regulation of oxidative stress pathway. As the most abundant peroxisomal enzyme, catalases catalyze decomposition of hydrogen peroxide [52]. To further study the mechanism of SntB mediated oxidative response of *A. flavus*, the *catC* (encode a catalase) gene was selected based on the above integration results. According to the peak map, the binding region of SntB on *catC* gene was significantly enriched in sntB-HA strains compared to WT strain (Figure 5F). The motif in the binding region was shown in Figure 5G and the sequence was TCCGCCCG. The relative expression levels of *catC* in WT and Δ*sntB* strains under MSB treatment were measured. As Figure 5H shown, the expression level of *catC* was significantly higher in Δ*sntB* strain than in WT strain, it suggested that to compensate the absence of *sntB*, *catC* is up-regulated to respond the higher intracellular oxidative level. However, under the stress of oxidant MSB, the deletion of *sntB* obvious suppressed the expression level of *catC* compared to that of WT strain, which reflected that the absence of *sntB* significantly impaired the capacity of *catC* to further respond to extra external oxidative stress. These results revealed that SntB is deeply involved in CatC mediated oxidative response in *A. flavus*.

**Figure 5.**
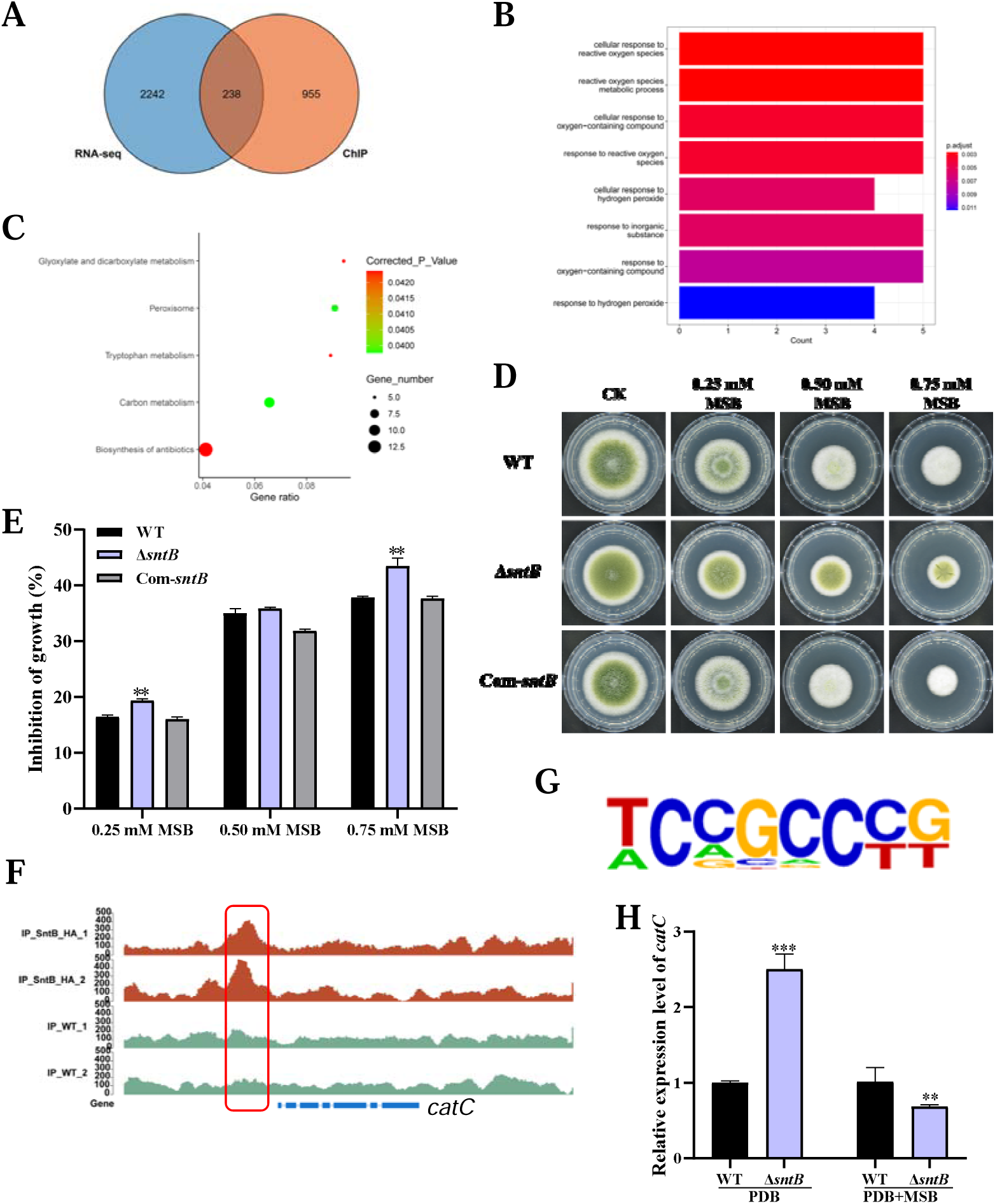
Integration of the results of ChIP-seq and RNA-seq assays. (A) Venn diagrams of ChIP-seq and RNA-seq. (B) GO analyses of the common genes. (C) KEGG analyses of the common genes. (D) The phenotype of WT, Δ*sntB*, and Com-*sntB* strains cultured in PDA containing a series concentration of MSB for 3 d. (E) Statistical analysis of the growth inhibition rate of MSB to all the above fungal strains according to Panel D. (F) Comparation of the enrich levels of the SntB binding region of *catC* gene between WT and sntB-HA strains. (G) The motif logo in the SntB binding region of *catC* gene. (H) The relative expression level of *catC* in WT and Δ*sntB* strains with or without MSB treatment.

### CatC is important for *A. flavus* response to oxidative stress

The functions of *catC* gene in *A. flavus* were further explored by knockout of the *catC* (Figure S5). As shown in Figure 6A-6C, the diameter of Δ*catC* strain was significantly smaller than that of WT, and the conidia number in the Δ*catC* strain decreased significantly compared to that of WT. The sclerotia production of Δ*catC* strain was also significantly less than that of WT strain (Figure 6D and 6E). In view of CatC is involved in the oxidative response pathways of *A. flavus* (Figure 4), both Δ*catC* and WT strains were treated by a serial concentration of MSB, and the results showed that the inhibition rates of MSB in Δ*catC* strain was significantly lower than that of WT (Figure 6F and 6G). Catalase is a major peroxisome protein and plays a critical role in removing peroxisome-generated ROS. The result of fluorescence intensity of oxidant-sensitive probe dichlorodihydrofluorescein-diacetate (DCFH-DA) showed that ROS accumulation in the Δ*catC* strain was higher than that in the WT strain (Figure 6G). This result echoed that the deletion of *sntB* increased intracellular oxidative level and the inhibition rate of MSB, and up-regulated the expression of *catC* (Figure 5D-5F). The role of CatC in the bio-synthesis of AFB1 was also assessed (Figure 6H). The results showed that a relatively large amount of AFB1 was produced by the Δ*catC* strain compared to the WT. But when under the stress of MSB, AFB1 yield of the WT strain was significantly more than that of Δ*catC* strain. All the above results revealed that the *catC* plays an important role in SntB mediated regulation pathway on fungal morphogenesis, oxidative stress responding, and AFB1 production.

**Figure 6.**
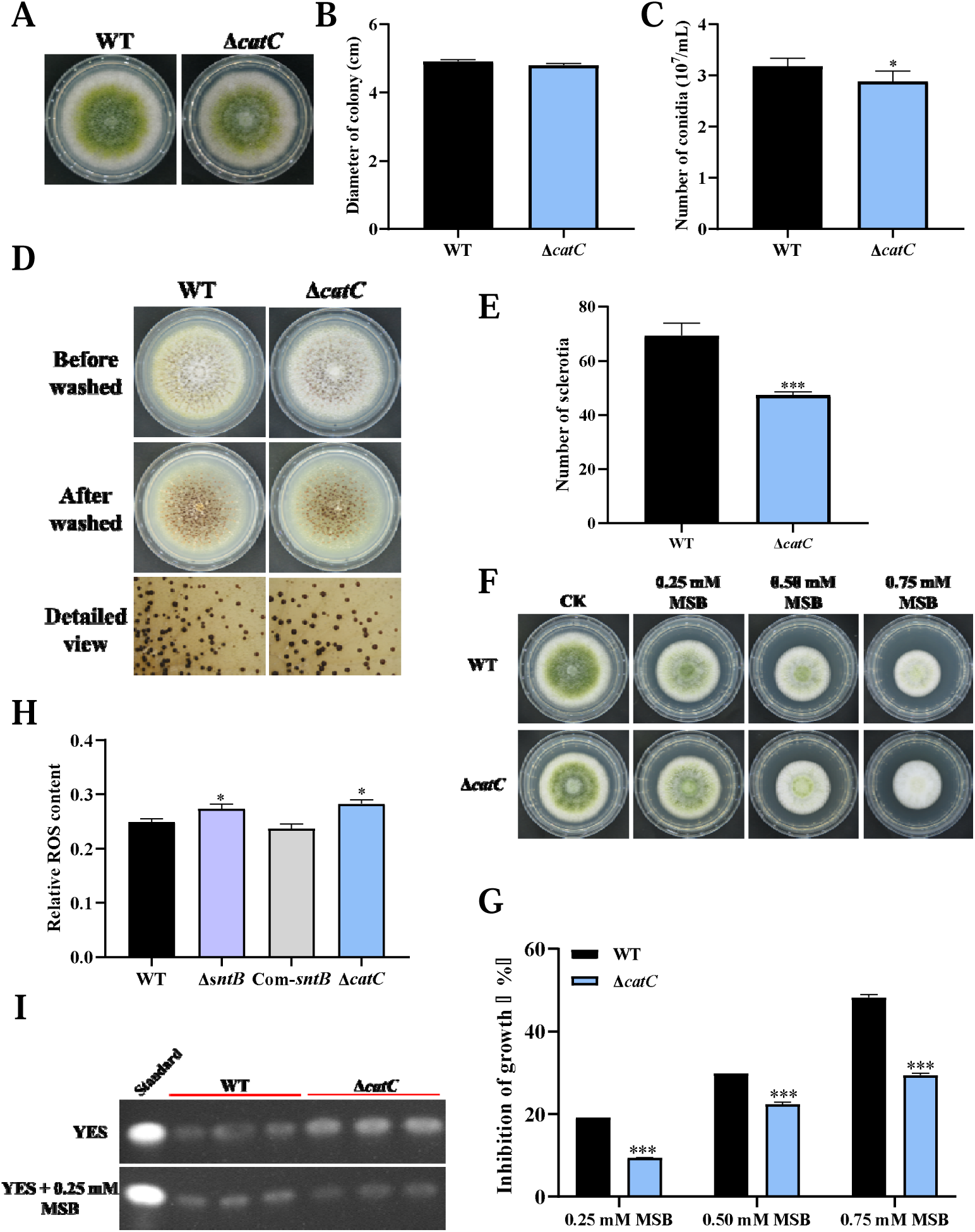
The functions of *catC* in *A. flavus*. (A) The colonies of WT and Δ*catC* strains grown on PDA at 37°C in dark for 4 d. (B) The colony diameter statistics of the above fungal strains. (C) The spore production statistics of the above fungal strains. (D) All above fungal strains were point-inoculated on CM medium and grown for 7 d at 37L. (E) The number of sclerotia of the above fungal strains. (F) The phenotype of above strains cultured in PDA medium containing a series concentration of MSB for 3 d. (G) Statistical analysis of the growth inhibition rate of MSB to all the above fungal strains according to (F). (H) Relative ROS level in the WT, Δ*sntB*, Com-sntB, and Δ*catC* strains. (I) AFB1 production of the above fungal strains was detected by TLC after the strains incubating at 29L in PDB medium for 7 d.

### SntB regulates fungal virulence through peroxidase mediated lipolysis

Biogenesis of peroxisomes was reported to promote lipid hydrolysis, produce the production of glycerol, and further change fungal pathogenicity [53]. Due to deeply involved in oxidative response of *A. flavus*, we wondered if SntB also takes part in the regulation of the production of lipid and glycerol. As shown in Table S14, one gene (G4B84_008359) in lipase activity GO term was significantly down-regulated in Δ*sntB* strain, which encoded a secretory lipase belonged to the virulence factors reported in *Pseudomonas aeruginosa* [54]. The lipase activity was also assayed by examining the ability to cleave glycerol tributyrate substrate [55]. The results showed that the colony diameter of Δ*sntB* strain on PDA medium with tributyrin were significantly smaller than that of the control, and the colony diameters of WT and Com-*sntB* strains on PDA medium were obviously bigger than those on 0.3% tributyrin PDA medium. The relative inhibition rate of tributyrin to the colony growth of Δ*sntB* strains was significantly higher than that of WT and Com-*sntB* strain (Figure S3D and S3E). Our previous study revealed that H3 lysine 36 trimethylation (H3K36me3) modification on the chromatin region of the sntB is regulated by AshA and SetB [34]. H3K36me3 usually promote gene transcription [56, 57]. Our study revealed the potential machinery associated with SntB mediated regulation on fungal morphogenesis, mycotoxin anabolism and fungal virulence, which lurks the axle of from SntB to fungal virulence and mycotoxin bio-synthesis through lipid catabolism (i.e. H3K36me3 modification-SntB-Peroxisomes-Lipid hydrolysis-fungal virulence and mycotoxin bio-synthesis).

## Discussion

SntB is a conserved regulator in many species, including *A. nidulans* [25], *Saccharomyces cerevisiae* [27], *Schizosaccharomyces pombe* [58], *F. oxysporum* [31], *Neurospora crassa* [32], and *A. flavus* [20, 25]. SntB can regulate the production of uncharacterized secondary metabolites, including aspergillicin A1 and aspergillicin F2 [59]. MoSntB protein was required for regulation of infection-associated autophagy in *M. oryzae* [51]. In *A. flavus*, the functions of *sntB* gene were previously analyzed by both Δ*sntB* and overexpression *sntB* genetic mutants[20]. SntB deletion impaired several developmental processes, such as sclerotia formation and heterokaryon compatibility, secondary metabolite synthesis, and ability to colonize host seeds, which were consistent with our results (Figure 1 and 2). Unlike, a complementation strain was constructed in this study which further clarified and confirmed the function of *sntB* gene. In this study, the potential mechanism under these effects were further analyzed by detection the related transcriptional factor genes of sporulation (*steA*, *WetA*, *fluG*, and *veA*), sclerotia formation (*nsdC*, *nsdD*, and *sclR*), and the AFs synthesis related genes *aflC*, *aflD*, *aflO*, *aflP*, and *aflR* (Figure S2). In the RNA-seq data, we also found some DEGs related to AFs synthesis (*aflB*, *aflE*, *aflH*, *aflK*, *aflN*, *aflO*, *aflP*, *aflQ*, *aflR*, *aflS*, *aflV*, and *aflW*) (Figure S6). And all these genes were down-regulated, which was consistent with that the AFs production in Δ*sntB* was significantly decreased compared to WT and Com-*sntB* (Figure 1F). These results inferred that SntB regulated the morphogenesis and the production of *A. flavus* through the above canonical signal pathways.

For the process of *A. flavus* invading hosts, in view of it is a notorious pathogen for plant and animal, we established both crop and insect models, especially silkworm represented animal mode profoundly revealed the critical role of SntB in fungal virulence (Figure 2). The results of crop kernel models showed that the number of spores of Δ*sntB* on kernels of both peanut and maize was dramatically lower than that of strains WT and Com-*sntB* (Figure 2A and 2B) and almost no AFB1 was detected on maize and peanut kernels inoculated with Δ*sntB*, while plenty of AFB1 were detected from the kernels infected by WT fungal strain, and AFB1 biosynthesis capacity of Com-*sntB* strains recovered compared to the Δ*sntB* and WT fungal strains (Figure 2C). These results were corroborated by previous study [20]. It was also found in this study that the survival rate of silkworms injected by spores of Δ*sntB* strain was significantly higher than the silkworms injected with spores from WT and Com-*sntB* fungal strains (Figure 2D and 2E). What’s more, we also assayed the effect of SntB on the activity amylase, which was closely related to the capacity of fungal infection [60]. As shown in Figure S3C, after adding iodine solution, the Δ*sntB* strain almost did not produce a degradation transparent ring compared with wild-type and complementary strains, indicating the amylase activity of the *sntB* gene knockout strain was significantly decreased (*P*<0.001). Our results comprehensively reveal the important function of SntB in the growth, development, secondary metabolite synthesis, and virulence of *A. flavus*.

SntB was reported as an important epigenetic reader. In *A. flavus*, SntB was reported to regulate global histone modifications (acetylation and methylation) and interact with EcoA and RpdA to form a conserved chromatin regulatory complex [20, 29]. Loss of *sntB* in *M. orzyae* also led to an increase in H3 acetylation [51]. In our RNA-seq data, we also found a set domain containing histone-lysine N-methyltransferase (Ash1, G4B84_009862) was down-regulated in Δ*sntB* strain compared to WT (Table S3), which was reported to regulate mycotoxin metabolism and virulence via H3K36 methylation in *A. flavus* [34]. Besides, SntB is reported to be a transcriptional regulator in *A. nidulans* [25] and *F. oxysporum* [31]. So, we use RNA-seq and ChIP-seq to study the transcriptional response of *sntB* in *A. flavus*. By integration analysis the results of ChIP-seq and RNA-seq assays, we found that the enriched GO terms and KEGG pathways of the DEGs were related to oxidative response (Figure 5A-5C). These results reflected that SntB plays an important role in fungal response to oxidative stress, which is consistent with previous reports that SntB could response to oxidative stress in yeast [27], *F. oxysporum* [31], and *M. oryzae* [51].

As a harmful by-product of oxidative metabolism, ROS is unavoidable and essential for fungus development [61, 62]. ROS has also been shown to be required for aflatoxin production [63–65]. Several oxidative stress-responsive transcription factors have been identified as regulating aflatoxin production, including AtfB, AP-1, and VeA [64, 66–68]. Previous studies shown that SntB protein coordinates the transcriptional response to hydrogen peroxide-mediated oxidative stress in the yeast [26, 27] and is involved in fungal respiration and ROS in *F. oxysporum* and *Neurospora crassa* [31, 32]. Several GO terms (“cellular response to reactive oxygen species”, “reactive oxygen species metabolic process”, and “cellular response to oxygen-containing compound”) and KEGG pathways (peroxisome) were enriched by the DEGs screened out by integration of ChIP-seq and RNA-seq data in this study (Figure 5). And the intracellular ROS level in the Δ*sntB* and Δ*catC* strains were significantly higher than that in WT strain (Figure 6H), which was similar to previous report on the *cat1* gene in *A. flavus* [69]. This is the first time to show that the SntB in *A. flavus* is important in oxidative stress response, through which SntB participates the regulating of aflatoxin bio-synthesis and fungal development.

Fungal defense against ROS is mediated by superoxide dismutases (SOD), catalases (CAT), and glutathione peroxidases (GPX). The effect of MSB on cellular growth and antioxidant enzyme induction in *A. flavus* was previously explored [70–72]. Once in the cell, menadione may release superoxide anion [73], which was scavenged by superoxide dismutases (SOD) and transformed into hydrogen peroxide, or react with nitric oxide to form peroxynitrite [74]. This study found that after knock out *sntB* gene, the strain growth was significantly inhibited by MSB (Figure 5F and 5G). Some genes encoded superoxide dismutases and catalases were reported to associated with AF/ST synthesis [75], including *mnSOD*, *sod1*, *sod2*, *catA*, *catB*, and *hyr1*. In our RNA-seq data, 7 related genes were screened out (Table S15), among which, bZIP transcription factor *Atf21* (G4B84_008675), *catlase C* (G4B84_009242), *catlase A* (G4B84_010740), superoxide dismutases *sod2* (G4B84_003204), peroxisomal membrane protein *PmpP24* (G4B84_001452) were up-regulated, while *catlase B* (G4B84_008381) and superoxide dismutase *sod1* (G4B84_009129) were down-regulated in Δ*sntB* strain, respectively. The binding region of SntB on *catA*, *catB*, *sod1*, *sod2*, and *catC* genes promoter were significantly enriched in sntB-HA strain compared to WT strain (Figure 5F and Figure S7A). Also, the motif in the binding regions were shown in Figure 5G and Figure S7B. These findings suggest that SNTB has the ability to interact with the promoter region of genes associated with oxidative processes, thereby modulating the expression of these genes.

Some studies reported the correlation between ROS formation, aflatoxin production, and antioxidant enzyme activation. Aflatoxin B1 biosynthesis and the activity of total superoxide dismutase were effectively inhibited by cinnamaldehyde, whereas the activities of catalase and glutathione peroxidase were opposite [76]. The expression of *catA*, *cat2* and *sod1*, and CAT enzymatic activity were opposite correlated to AFB1 biosynthesis under AFs inhibitor piperine treatment [77]. Deletion the gene of *sod* (GenBank accession no: CA747446) reduced AFs production [78], which was most similar to *sod2* (G4B84_003204). The mitochondria-specific *sod* and the genes *aflA*, *aflM*, and *aflP* belonging to the AFs gene cluster were reported to co-regulated [79]. Ethanol can inhibit fungal growth and AFB_1_ production in *A. flavus* and enhanced levels of anti-oxidant enzymatic genes, including *Cat*, *Cat1*, *Cat2*, *CatA*, and Cu, Zn superoxide dismutase gene *sod1*. All these reports indicated that the expression of anti-oxidant enzymatic genes was opposite correlated to AFB1 biosynthesis.

In our study, 7 genes related to oxidative response were obviously differentially expressed in transcriptome data (Table S15). Among these DEGs, 5 out of 7 genes were up-regulated in Δ*sntB* strain. Based on the AFs production in Δ*sntB* was significantly decreased compared to WT and Com-*sntB* (Figure 1F), the most up-regulated gene in Δ*sntB* strain, *catC* (G4B84_000242), was selected for further analysis. We found that the deletion of *sntB* significantly up-regulated the *catC* gene, however, the expression of the *catC* gene was almost undetectable under MSB treatment (Figure 5F). Results also showed that the inhibition rates of MSB to Δ*catC* strain was significantly lower than that of WT group and AFB1 yield of the Δ*catC* strain was significantly decreased than that of WT strain under the stress of MSB (Figure 6F-6H). These results indicated that SntB is profoundly involved in the *catC* mediated oxidative response.

Peroxisomes are intimately associated with the metabolism of lipid droplets [80] and the histone lysine methyltransferase ASH1 promotes peroxisome biogenesis, inhibits lipolysis, and further affects pathogenesis of *Metarhizium robertsii* [53]. Set2 histone methyltransferase family in *A. flavus*, AshA and SetB, were found to regulate mycotoxin metabolism and virulence via H3K36me3, including the chromatin region of the *sntB* [34]. By ChIP-seq and RNA-seq, SntB was found to essential for *A. flavus* to maintain the homeostasis of intracellular reactive oxygen species (Figure 5A) and several anti-oxidant enzymes were up-regulated in Δ*sntB* strain (Table S14). In addition, we also found only one down-regulated DEG (G4B84_008359) in lipase activity GO term in our RNA-seq data (Table S14), which encodes a secretory lipase and belongs to the virulence factors reported in *Pseudomonas aeruginosa* [54]. These results suggested that SntB plays a pivotal role in regulating peroxisome biogenesis to promote lipolysis involving in fungal pathogenesis.

Overall, we explored and clarified the bio-function of the SntB and found that SntB responses to oxidative stress through related oxidoreductase represented by CatC in *A. flavus* (Figure 7). Our study revealed the potential machinery associated with SntB mediated regulation on fungal morphogenesis, mycotoxin anabolism, and fungal virulence, which lurks the axle of from SntB to fungal virulence and mycotoxin bio-synthesis (i.e. SntB-Peroxisomes-Lipid hydrolysis-fungal virulence and mycotoxin bio-synthesis). The work of this study provided a novel perspective for developing new prevention and control strategies against pathogenic fungi.

**Figure 7.**
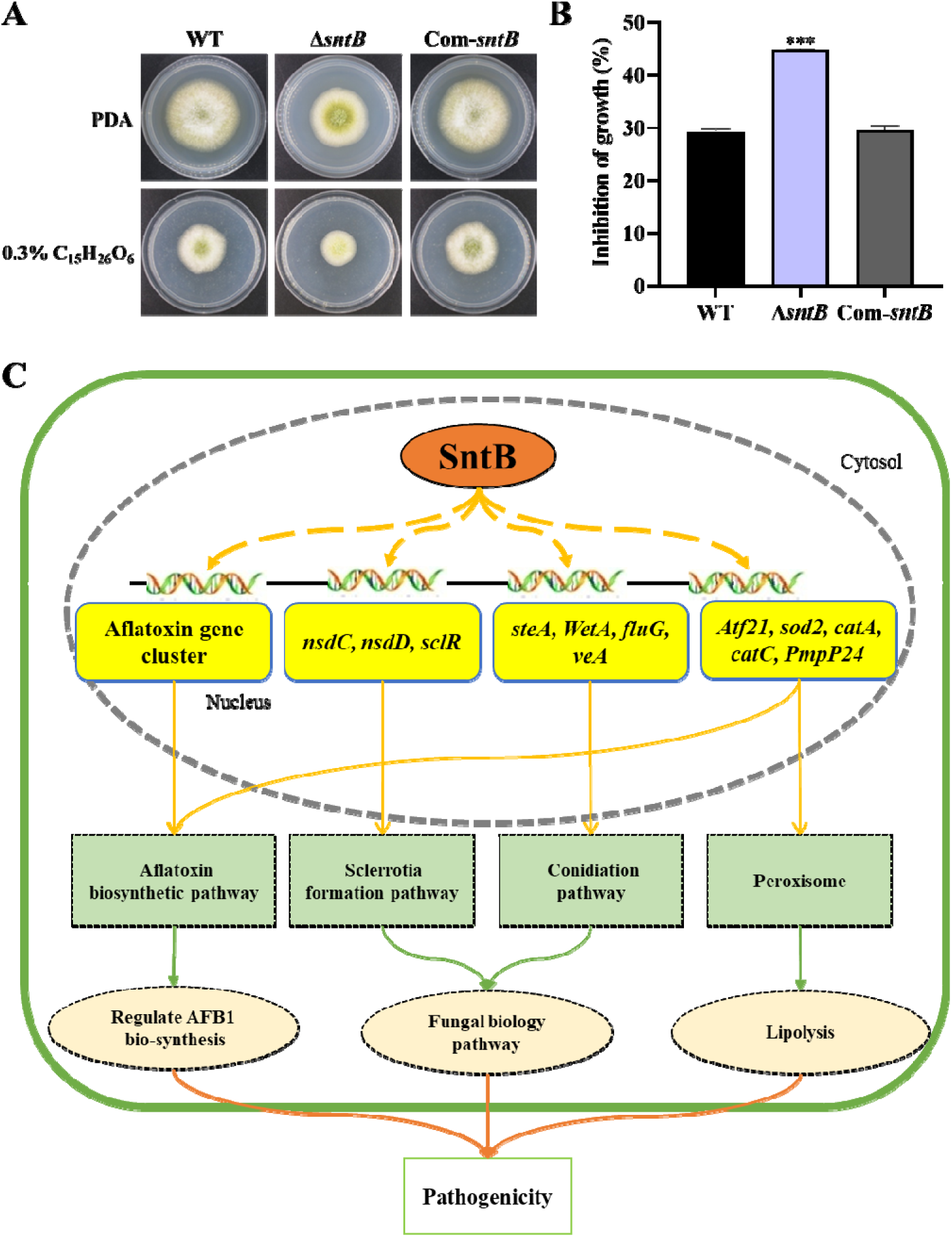
SntB regulate peroxisome biogenesis, fatty acid utilization, and fungal pathogenicity in *A. flavus*. (A) The phenotype of each strain on PDA medium containing 0.3% tributyrin, (B) Statistics of inhibition rates. The asterisk *** above the bars represents significantly different (*p*<0.001). (C) Mechanistic diagram of the bio-functions of SntB in *A. flavus*.

## Compliance with ethical requirements

This article does not contain any studies with human or animal subjects.

## Data Availability

All data needed to evaluate the conclusions are present in the paper and/or the Supporting Information. Raw data of the ChIP and RNA-seq were submitted to GSE247683.

## Acknowledgement

We especially thank Professor Jun Yuan, Xiuna Wang, Yu Wang and Xinyi Nie for their support in instrument maintenance and reagent ordering.

## Supplementary figures

**Figure S1.**
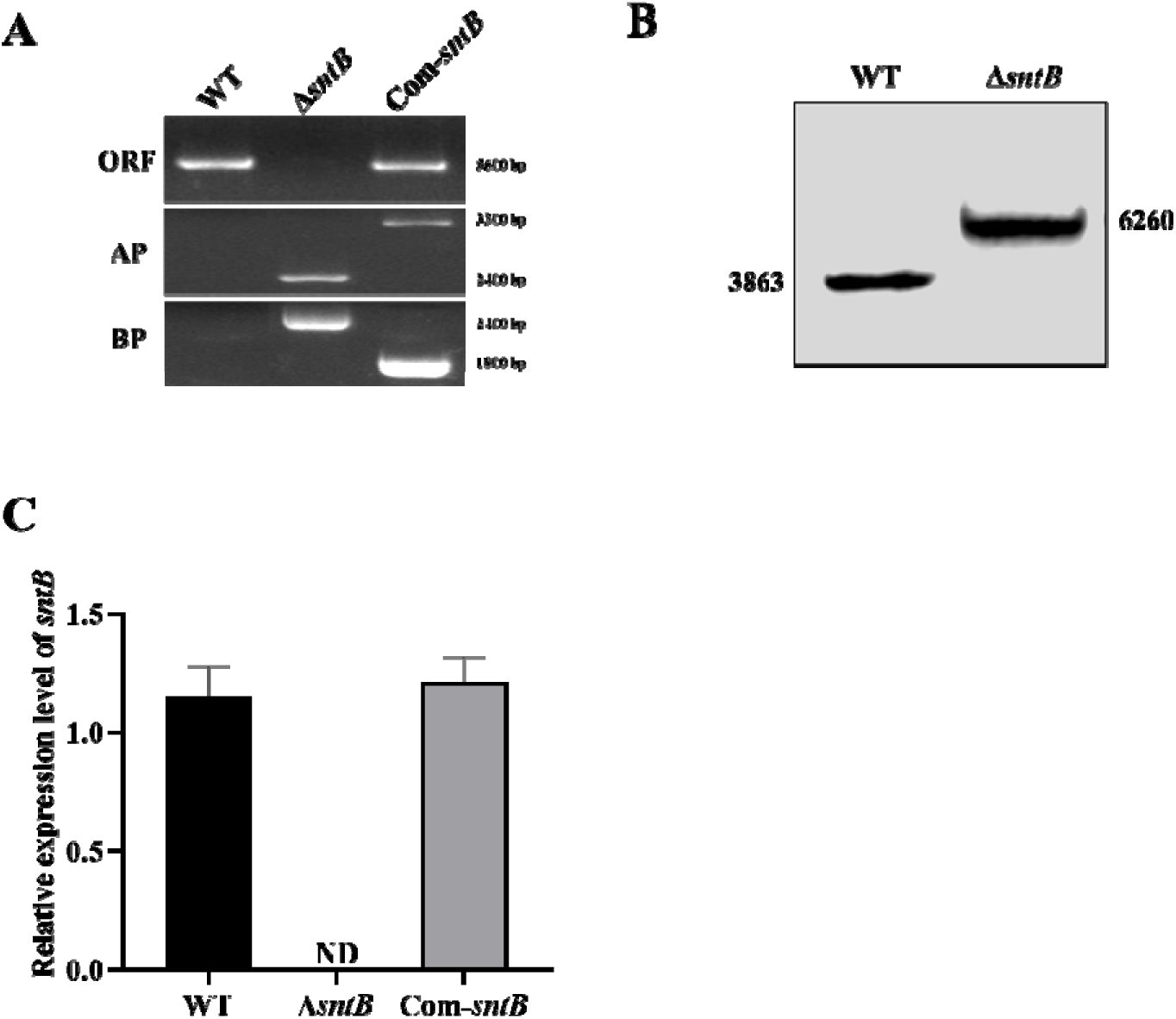
The construction of mutant strains. (A) PCR verification of gDNA in WT, Δ*sntB* and Com-*sntB* strains (“ORF” represents the sntB gene fragment, “AP” represents the amplification of the fusion fragment upstream with primers *sntB*-p1 and P801, and “BP” represents the downstream of the fusion fragment from primers P1020 and *sntB*-p4). (B) Southern blot analysis result of WT, Δ*sntB* candidate strains (The genome DNA of each strain was digested with restrictive endonuclease EcoR I and hybridized with the probe of 1, 533 bp). (C) qRT-PCR verification of the expression level of the *sntB* gene in WT and *sntB* gene mutant strains.

**Figure S2.**
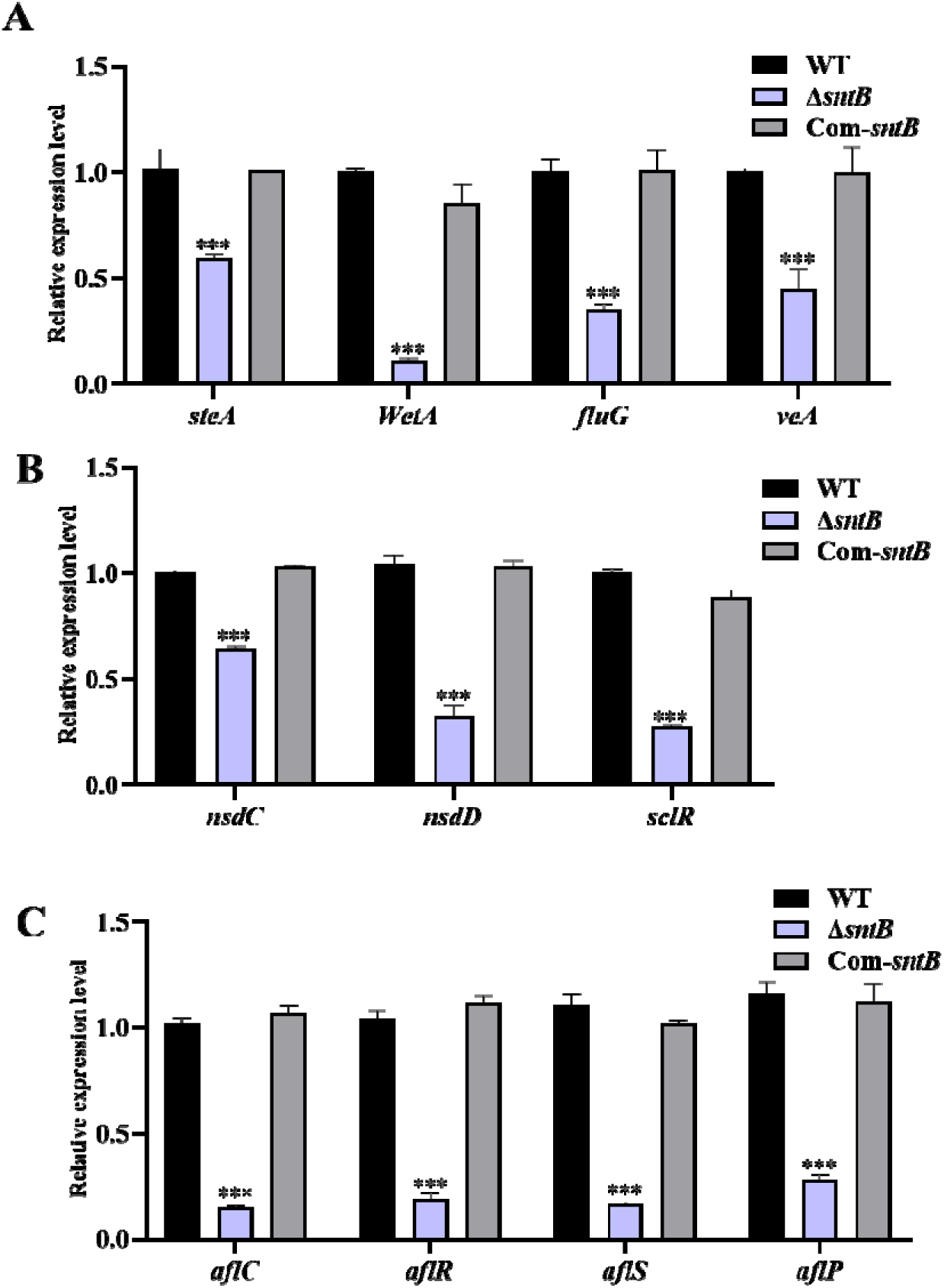
The expression of genes related to sporulation, sclerotia production, and aflatoxin synthesis. (A) The expression of sporulation-related genes *steA*, *WetA*, *fluG*, and *veA* in each strain at 48 h. (B) The expression of sclerotia-associated genes *nsdC*, *nsdD*, and *sclR* in each strain at 48 h. (C) The expression of aflatoxin-associated genes in each strain at 48 h. The asterisk *** above the bars represents significantly different (p<0.001)

**Figure S3.**
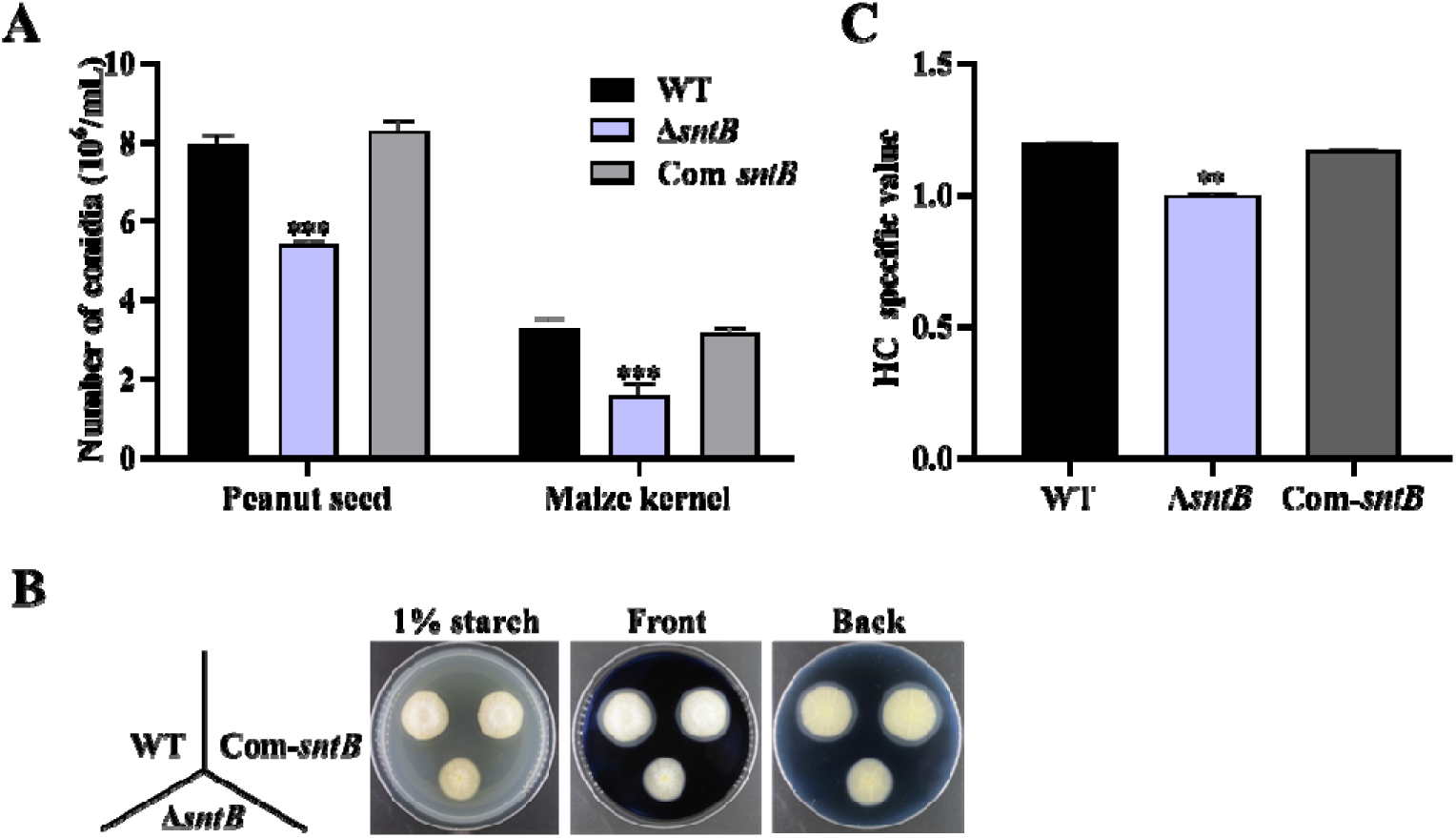
The changes of number of conidia, amylase, and lipase in WT, Δ*sntB*, and Com-*sntB* strains. (A) Statistics of the number of conidia on the corn seed. (B) The phenotype of each strain on starch screening medium supplemented with 0.1% of soluble starch in darkness at 29℃ for 3 d, followed by addition of iodine solution. (C) HC value of the clear circle (Outer diameter/inner diameter) of the salient analysis of the map. The asterisk ** above the bars represents significantly different (*p*<0.01).

**Figure S4.**
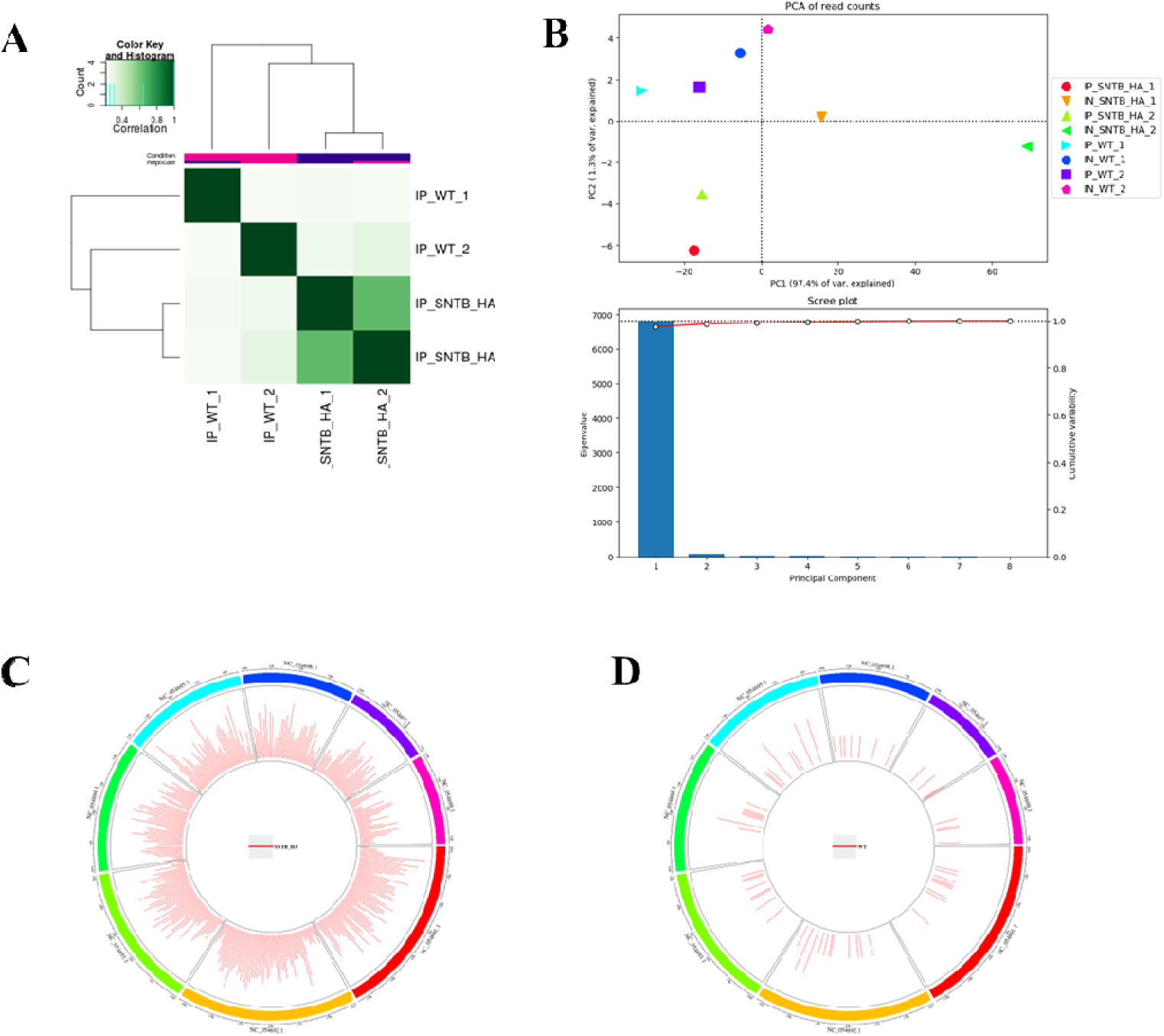
Sequence information of ChIP-seq. (A) Heatmap. (B) PCA. (C) Peak distribution on the genome of SNTB-HA group. (D) Peak distribution on the genome of WT group.

**Figure S5.**
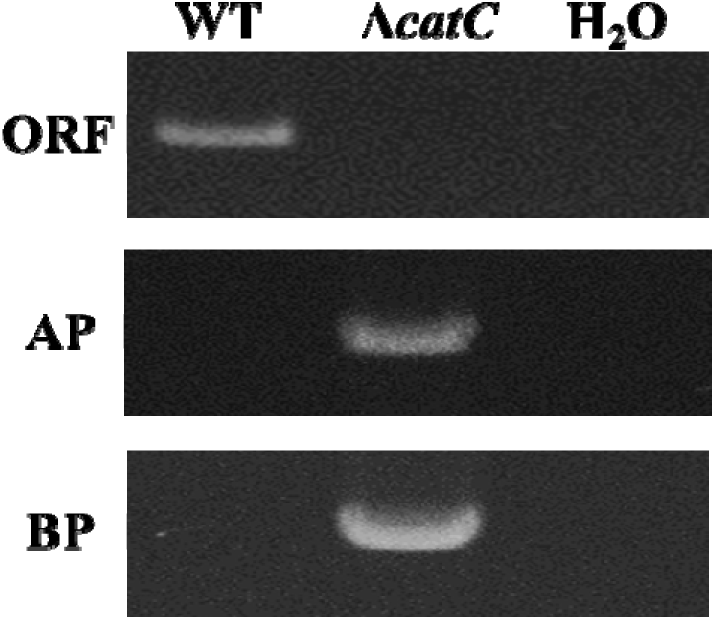
PCR verification of gDNA in WT and Δ*catC*

**Figure S6.**
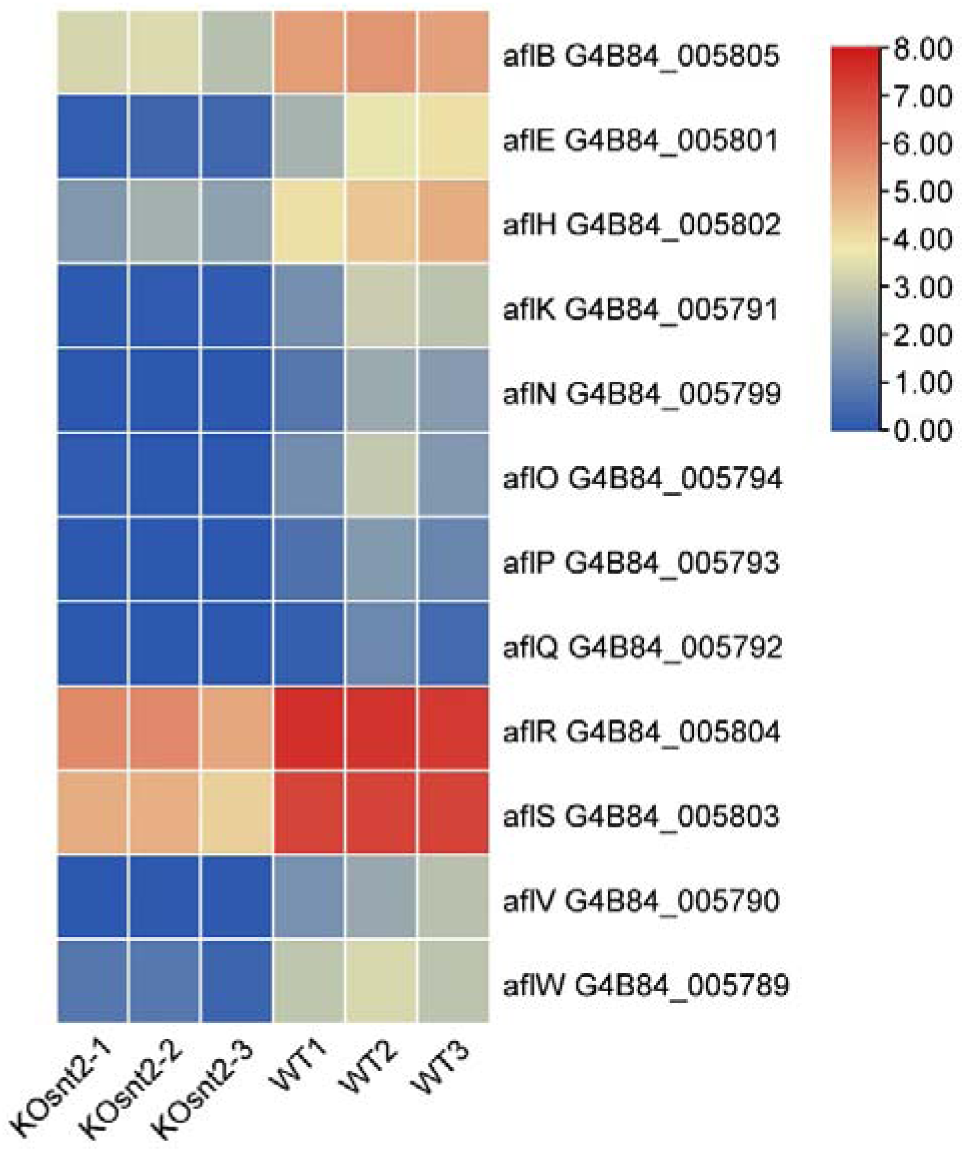
Heatmap of the DEGs related to oxidative response in transcriptome data draw by TBtools.

**Figure S7.**
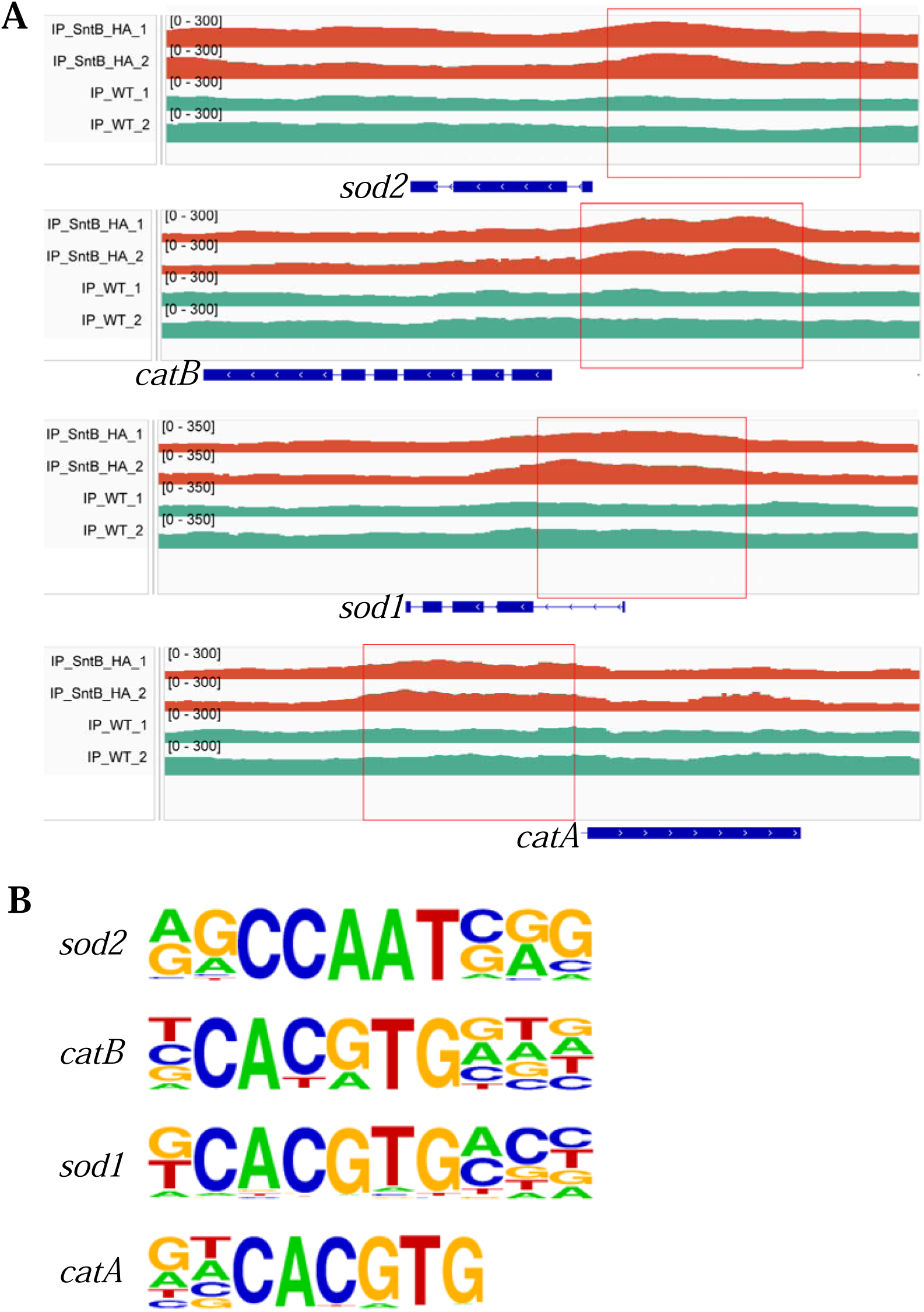
The binding region and motif of SntB on the *catA*, *catB*, *sod1*, and *sod2* genes. (A) Comparation of the enrich levels of the SntB binding region of *catA*, *catB*, *sod1*, and *sod2* genes between WT and sntB-HA strains. (B) The motif logo in the SntB binding region of *catA*, *catB*, *sod1*, and *sod2* genes.

## Supplementary table captions

Table S1. Sequencing data statistics in RNA-seq.

Table S2. Alignment results of each sample in RNA-seq.

Table S3. The information of DEGs in transcriptome data.

Table S4. GO enriched in the DEGs in transcriptome data.

Table S5. KEGG pathways enriched of the DEGs in transcriptome data.

Table S6. Sequencing data statistics in ChIP-seq.

Table S7. Alignment results of each sample in ChIP-seq.

Table S8. The information of up regulated peak in ChIP-seq data.

Table S9. Known motifs identified by HOMER motif enrichment analysis software.

Table S10. GO enriched in the up regulated genes in ChIP-seq data.

Table S11. KEGG enriched in the up regulated genes in ChIP-seq data.

Table S12. GO enriched in the 238 common DEGs in RNA-seq and ChIP-seq data.

Table S13. KEGG enriched in the 238 common DEGs in RNA-seq and ChIP-seq data.

Table S14. The information of DEGs related to lipase activity in transcriptome data.

Table S15. The information of DEGs related to oxidative response in transcriptome data.

## References

[1] Hedayati M.T., Pasqualotto A.C., Warn P.A., Bowyer P., Denning D.W. Aspergillus flavus: human pathogen, allergen and mycotoxin producer [J]. Microbiology (Reading), 2007, 153(Pt 6): 1677–1692.

[2] Guan Y., Chen J., Nepovimova E., Long M., Wu W., Kuca K. Aflatoxin Detoxification Using Microorganisms and Enzymes [J]. Toxins (Basel), 2021, 13(1).

[3] Mishra H.N., Das C. A review on biological control and metabolism of aflatoxin [J]. Crit Rev Food Sci Nutr, 2003, 43(3): 245–264.

[4] Vardon P.J., Mclaughlin C., Nardinelli C. Potential economic costs of mycotoxins in the United States [J]. 2003.

[5] Riba A., Bouras N., Mokrane S., Mathieu F., Lebrihi A., Sabaou N. Aspergillus section Flavi and aflatoxins in Algerian wheat and derived products [J]. Food Chem Toxicol, 2010, 48(10): 2772–2777.

[6] Plaz Torres M.C., Bodini G., Furnari M., Marabotto E., Zentilin P., Giannini E.G. Nuts and Non-Alcoholic Fatty Liver Disease: Are Nuts Safe for Patients with Fatty Liver Disease? [J]. Nutrients, 2020, 12(11).

[7] Cleveland T.E., Yu J., Fedorova N., Bhatnagar D., Payne G.A., Nierman W.C., Bennett J.W. Potential of Aspergillus flavus genomics for applications in biotechnology [J]. Trends Biotechnol, 2009, 27(3): 151–157.

[8] Yabe K., Nakajima H. Enzyme reactions and genes in aflatoxin biosynthesis [J]. Appl Microbiol Biotechnol, 2004, 64(6): 745–755.

[9] Yu J. Current understanding on aflatoxin biosynthesis and future perspective in reducing aflatoxin contamination [J]. Toxins (Basel), 2012, 4(11): 1024–1057.

[10] Roze L.V., Hong S.Y., Linz J.E. Aflatoxin biosynthesis: current frontiers [J]. Annu Rev Food Sci Technol, 2013, 4: 293–311.

[11] Ehrlich K.C. Predicted roles of the uncharacterized clustered genes in aflatoxin biosynthesis [J]. Toxins (Basel), 2009, 1(1): 37–58.

[12] Chang P.K. The Aspergillus parasiticus protein AFLJ interacts with the aflatoxin pathway-specific regulator AFLR [J]. Mol Genet Genomics, 2003, 268(6): 711–719.

[13] Price M.S., Yu J., Nierman W.C., Kim H.S., Pritchard B., Jacobus C.A., Bhatnagar D., Cleveland T.E., Payne G.A. The aflatoxin pathway regulator AflR induces gene transcription inside and outside of the aflatoxin biosynthetic cluster [J]. FEMS Microbiol Lett, 2006, 255(2): 275–279.

[14] Xu J., Luo X. Molecular biology of aflatoxin biosynthesis [J]. Wei Sheng Yan Jiu, 2003, 32(6): 628–631.

[15] Yu J.H., Keller N. Regulation of secondary metabolism in filamentous fungi [J]. Annu Rev Phytopathol, 2005, 43: 437–458.

[16] Georgianna D.R., Payne G.A. Genetic regulation of aflatoxin biosynthesis: from gene to genome [J]. Fungal Genet Biol, 2009, 46(2): 113–125.

[17] Montibus M., Pinson-Gadais L., Richard-Forget F., Barreau C., Ponts N. Coupling of transcriptional response to oxidative stress and secondary metabolism regulation in filamentous fungi [J]. Crit Rev Microbiol, 2015, 41(3): 295–308.

[18] Lv Y., Wang J., Yang H., Li N., Farzaneh M., Wei S., Zhai H., Zhang S., Hu Y. Lysine 2-hydroxyisobutyrylation orchestrates cell development and aflatoxin biosynthesis in Aspergillus flavus [J]. Environ Microbiol, 2022, 24(9): 4356–4368.

[19] Yang G., Yue Y., Ren S., Yang M., Zhang Y., Cao X., Wang Y., Zhang J., Ge F., Wang S. Lysine acetylation contributes to development, aflatoxin biosynthesis and pathogenicity in Aspergillus flavus [J]. Environ Microbiol, 2019, 21(12): 4792–4807.

[20] Pfannenstiel B.T., Greco C., Sukowaty A.T., Keller N.P. The epigenetic reader SntB regulates secondary metabolism, development and global histone modifications in Aspergillus flavus [J]. Fungal Genet Biol, 2018, 120: 9–18.

[21] Ren S., Yang M., Yue Y., Ge F., Li Y., Guo X., Zhang J., Zhang F., Nie X., Wang S. Lysine Succinylation Contributes to Aflatoxin Production and Pathogenicity in Aspergillus flavus [J]. Mol Cell Proteomics, 2018, 17(3): 457–471.

[22] Wang Y., Yang M., Ge F., Jiang B., Hu R., Zhou X., Yang Y., Liu M. Lysine Succinylation of VBS Contributes to Sclerotia Development and Aflatoxin Biosynthesis in Aspergillus flavus [J]. Mol Cell Proteomics, 2023, 22(2): 100490.

[23] Lv Y. Proteome-wide profiling of protein lysine acetylation in Aspergillus flavus [J]. PLoS One, 2017, 12(6): e0178603.

[24] Wang J., Liang L., Wei S., Zhang S., Hu Y., Lv Y. Histone 2-Hydroxyisobutyryltransferase Encoded by Afngg1 Is Involved in Pathogenicity and Aflatoxin Biosynthesis in Aspergillus flavus [J]. Toxins (Basel), 2022, 15(1).

[25] Pfannenstiel B.T., Zhao X., Wortman J., Wiemann P., Throckmorton K., Spraker J.E., Soukup A.A., Luo X., Lindner D.L., Lim F.Y., Knox B.P., Haas B., Fischer G.J., Choera T., Butchko R.A.E., Bok J.W., Affeldt K.J., Keller N.P., Palmer J.M. Revitalization of a Forward Genetic Screen Identifies Three New Regulators of Fungal Secondary Metabolism in the Genus Aspergillus [J]. mBio, 2017, 8(5).

[26] Singh R.K., Gonzalez M., Kabbaj M.H., Gunjan A. Novel E3 ubiquitin ligases that regulate histone protein levels in the budding yeast Saccharomyces cerevisiae [J]. PLoS One, 2012, 7(5): e36295.

[27] Baker L.A., Ueberheide B.M., Dewell S., Chait B.T., Zheng D., Allis C.D. The yeast Snt2 protein coordinates the transcriptional response to hydrogen peroxide-mediated oxidative stress [J]. Mol Cell Biol, 2013, 33(19): 3735–3748.

[28] Tannous J., Barda O., Luciano-Rosario D., Prusky D.B., Sionov E., Keller N.P. New Insight Into Pathogenicity and Secondary Metabolism of the Plant Pathogen Penicillium expansum Through Deletion of the Epigenetic Reader SntB [J]. Front Microbiol, 2020, 11: 610.

[29] Karahoda B., Pardeshi L., Ulas M., Dong Z., Shirgaonkar N., Guo S., Wang F., Tan K., Sarikaya-Bayram Ö., Bauer I., Dowling P., Fleming A.B., Pfannenstiel B.T., Luciano-Rosario D., Berger H., Graessle S., Alhussain M.M., Strauss J., Keller N.P., Wong K.H., Bayram Ö. The KdmB-EcoA-RpdA-SntB chromatin complex binds regulatory genes and coordinates fungal development with mycotoxin synthesis [J]. Nucleic Acids Res, 2022, 50(17): 9797–9813.

[30] Karahoda B., Pfannenstiel B.T., Sarikaya-Bayram Ö., Dong Z., Ho Wong K., Fleming A.B., Keller N.P., Bayram Ö. The KdmB-EcoA-RpdA-SntB (KERS) chromatin regulatory complex controls development, secondary metabolism and pathogenicity in Aspergillus flavus [J]. Fungal Genet Biol, 2023, 169: 103836.

[31] Denisov Y., Freeman S., Yarden O. Inactivation of Snt2, a BAH/PHD-containing transcription factor, impairs pathogenicity and increases autophagosome abundance in Fusarium oxysporum [J]. Mol Plant Pathol, 2011, 12(5): 449–461.

[32] Denisov Y., Yarden O., Freeman S. The transcription factor SNT2 is involved in fungal respiration and reactive oxidative stress in Fusarium oxysporum and Neurospora crassa [J]. Physiological and Molecular Plant Pathology, 2011, 76(2): 137–143.

[33] Chang P.K., Scharfenstein L.L., Wei Q., Bhatnagar D. Development and refinement of a high-efficiency gene-targeting system for Aspergillus flavus [J]. J Microbiol Methods, 2010, 81(3): 240–246.

[34] Zhuang Z., Pan X., Zhang M., Liu Y., Huang C., Li Y., Hao L., Wang S. Set2 family regulates mycotoxin metabolism and virulence via H3K36 methylation in pathogenic fungus Aspergillus flavus [J]. Virulence, 2022, 13(1): 1358–1378.

[35] Pan X., Hao L., Yang C., Lin H., Wu D., Chen X., Zhang M., Ma D., Wang Y., Fu W., Yao Y., Wang S., Zhuang Z. SWD1 epigenetically chords fungal morphogenesis, aflatoxin biosynthesis, metabolism, and virulence of Aspergillus flavus [J]. Journal of Hazardous Materials, 2023: 131542.

[36] Hu Y., Yang G., Zhang D., Liu Y., Li Y., Lin G., Guo Z., Wang S., Zhuang Z. The PHD Transcription Factor Rum1 Regulates Morphogenesis and Aflatoxin Biosynthesis in [J]. Toxins, 2018, 10(7).

[37] Pan X., Hao L., Yang C., Lin H., Wu D., Chen X., Zhang M., Ma D., Wang Y., Fu W., Yao Y., Wang S., Zhuang Z. SWD1 epigenetically chords fungal morphogenesis, aflatoxin biosynthesis, metabolism, and virulence of Aspergillus flavus [J]. J Hazard Mater, 2023, 455: 131542.

[38] Kaminskyj S.G. Septum position is marked at the tip of Aspergillus nidulans hyphae [J]. Fungal Genet Biol, 2000, 31(2): 105–113.

[39] Zhang Y., Fan J., Ye J., Lu L. The fungal-specific histone acetyltransferase Rtt109 regulates development, DNA damage response, and virulence in Aspergillus fumigatus [J]. Mol Microbiol, 2021, 115(6): 1191–1206.

[40] Wen M., Lan H., Sun R., Chen X., Zhang X., Zhu Z., Tan C., Yuan J., Wang S. Histone deacetylase SirE regulates development, DNA damage response and aflatoxin production in Aspergillus flavus [J]. Environmental Microbiology, 2022.

[41] Hao L., Zhang M., Yang C., Pan X., Wu D., Lin H., Ma D., Yao Y., Fu W., Chang J., Yang Y., Zhuang Z. The epigenetic regulator Set9 harmonizes fungal development, secondary metabolism, and colonization capacity of Aspergillus flavus [J]. International Journal of Food Microbiology, 2023, 403: 110298.

[42] Liao Y., Smyth G.K., Shi W. featureCounts: an efficient general purpose program for assigning sequence reads to genomic features [J]. Bioinformatics, 2013, 30(7): 923–930.

[43] Li H., Durbin R. Fast and accurate short read alignment with Burrows-Wheeler transform [J]. Bioinformatics, 2009, 25(14): 1754–1760.

[44] Hull R.P., Srivastava P.K., D’Souza Z., Atanur S.S., Mechta-Grigoriou F., Game L., Petretto E., Cook H.T., Aitman T.J., Behmoaras J. Combined ChIP-Seq and transcriptome analysis identifies AP-1/JunD as a primary regulator of oxidative stress and IL-1β synthesis in macrophages [J]. BMC Genomics, 2013, 14: 92.

[45] Zhou X., Su Z. EasyGO: Gene Ontology-based annotation and functional enrichment analysis tool for agronomical species [J]. BMC Genomics, 2007, 8: 246.

[46] Xie C., Mao X., Huang J., Ding Y., Wu J., Dong S., Kong L., Gao G., Li C.Y., Wei L. KOBAS 2.0: a web server for annotation and identification of enriched pathways and diseases [J]. Nucleic Acids Res, 2011, 39(Web Server issue): W316–322.

[47] Mengjuan Z., Guanglan L., Xiaohua P., Weitao S., Can T., Xuan C., Yanling Y., Zhenhong Z. The PHD transcription factor Cti6 is involved in the fungal colonization and aflatoxin B1 biological synthesis of Aspergillus flavus [J]. IMA Fungus, 2021, 12(1): 12.

[48] Hu Y., Yang G., Zhang D., Liu Y., Li Y., Lin G., Guo Z., Wang S., Zhuang Z. The PHD Transcription Factor Rum1 Regulates Morphogenesis and Aflatoxin Biosynthesis in Aspergillus flavus [J]. Toxins (Basel), 2018, 10(7): 301.

[49] Cary J.W., Harris-Coward P.Y., Ehrlich K.C., Mack B.M., Kale S.P., Larey C., Calvo A.M. NsdC and NsdD affect Aspergillus flavus morphogenesis and aflatoxin production [J]. Eukaryot Cell, 2012, 11(9): 1104–1111.

[50] Yang G., Hu Y., Fasoyin O.E., Yue Y., Chen L., Qiu Y., Wang X., Zhuang Z., Wang S. The Phosphatase CDC14 Regulates Development, Aflatoxin Biosynthesis and Pathogenicity [J]. Frontiers In Cellular and Infection Microbiology, 2018, 8: 141.

[51] He M., Xu Y., Chen J., Luo Y., Lv Y., Su J., Kershaw M.J., Li W., Wang J., Yin J., Zhu X., Liu X., Chern M., Ma B., Wang J., Qin P., Chen W., Wang Y., Wang W., Ren Z., Wu X., Li P., Li S., Peng Y., Lin F., Talbot N.J., Chen X. MoSnt2-dependent deacetylation of histone H3 mediates MoTor-dependent autophagy and plant infection by the rice blast fungus Magnaporthe oryzae [J]. Autophagy, 2018, 14(9): 1543–1561.

[52] Okumoto K., Tamura S., Honsho M., Fujiki Y. Peroxisome: Metabolic Functions and Biogenesis [J]. Adv Exp Med Biol, 2020, 1299: 3–17.

[53] Wang L., Lai Y., Chen J., Cao X., Zheng W., Dong L., Zheng Y., Li F., Wei G., Wang S. The ASH1-PEX16 regulatory pathway controls peroxisome biogenesis for appressorium-mediated insect infection by a fungal pathogen [J]. Proc Natl Acad Sci U S A, 2023, 120(4): e2217145120.

[54] Papadopoulos A., Busch M., Reiners J., Hachani E., Baeumers M., Berger J., Schmitt L., Jaeger K.E., Kovacic F., Smits S.H.J., Kedrov A. The periplasmic chaperone Skp prevents misfolding of the secretory lipase A from Pseudomonas aeruginosa [J]. Front Mol Biosci, 2022, 9: 1026724.

[55] Sieber M.H., Thummel C.S. The DHR96 nuclear receptor controls triacylglycerol homeostasis in Drosophila [J]. Cell Metab, 2009, 10(6): 481–490.

[56] Zhao W., Neyt P., Van Lijsebettens M., Shen W.H., Berr A. Interactive and noninteractive roles of histone H2B monoubiquitination and H3K36 methylation in the regulation of active gene transcription and control of plant growth and development [J]. New Phytol, 2019, 221(2): 1101–1116.

[57] Kooistra S.M., Helin K. Molecular mechanisms and potential functions of histone demethylases [J]. Nat Rev Mol Cell Biol, 2012, 13(5): 297–311.

[58] Roguev A., Shevchenko A., Schaft D., Thomas H., Stewart A.F., Shevchenko A. A comparative analysis of an orthologous proteomic environment in the yeasts Saccharomyces cerevisiae and Schizosaccharomyces pombe [J]. Mol Cell Proteomics, 2004, 3(2): 125–132.

[59] Greco C., Pfannenstiel B.T., Liu J.C., Keller N.P. Depsipeptide Aspergillicins Revealed by Chromatin Reader Protein Deletion [J]. ACS Chem Biol, 2019, 14(6): 1121–1128.

[60] Mellon J.E., Cotty P.J., Dowd M.K. Aspergillus flavus hydrolases: their roles in pathogenesis and substrate utilization [J]. Appl Microbiol Biotechnol, 2007, 77(3): 497–504.

[61] Aguirre J., Ríos-Momberg M., Hewitt D., Hansberg W. Reactive oxygen species and development in microbial eukaryotes [J]. Trends Microbiol, 2005, 13(3): 111–118.

[62] Gessler N.N., Aver’yanov A.A., Belozerskaya T.A. Reactive oxygen species in regulation of fungal development [J]. Biochemistry (Mosc), 2007, 72(10): 1091–1109.

[63] Jayashree T., Subramanyam C. Oxidative stress as a prerequisite for aflatoxin production by Aspergillus parasiticus [J]. Free Radic Biol Med, 2000, 29(10): 981–985.

[64] Roze L.V., Chanda A., Wee J., Awad D., Linz J.E. Stress-related transcription factor AtfB integrates secondary metabolism with oxidative stress response in aspergilli [J]. J Biol Chem, 2011, 286(40): 35137–35148.

[65] Yang L., Fountain J.C., Wang H., Ni X., Ji P., Lee R.D., Kemerait R.C., Scully B.T., Guo B. Stress Sensitivity Is Associated with Differential Accumulation of Reactive Oxygen and Nitrogen Species in Maize Genotypes with Contrasting Levels of Drought Tolerance [J]. Int J Mol Sci, 2015, 16(10): 24791–24819.

[66] Reverberi M., Zjalic S., Ricelli A., Punelli F., Camera E., Fabbri C., Picardo M., Fanelli C., Fabbri A.A. Modulation of antioxidant defense in Aspergillus parasiticus is involved in aflatoxin biosynthesis: a role for the ApyapA gene [J]. Eukaryot Cell, 2008, 7(6): 988–1000.

[67] Sakamoto K., Arima T.H., Iwashita K., Yamada O., Gomi K., Akita O. Aspergillus oryzae atfB encodes a transcription factor required for stress tolerance in conidia [J]. Fungal Genet Biol, 2008, 45(6): 922–932.

[68] Baidya S., Duran R.M., Lohmar J.M., Harris-Coward P.Y., Cary J.W., Hong S.Y., Roze L.V., Linz J.E., Calvo A.M. VeA is associated with the response to oxidative stress in the aflatoxin producer Aspergillus flavus [J]. Eukaryot Cell, 2014, 13(8): 1095–1103.

[69] Zhu Z., Yang M., Bai Y., Ge F., Wang S. Antioxidant-related catalase CTA1 regulates development, aflatoxin biosynthesis, and virulence in pathogenic fungus Aspergillus flavus [J]. Environ Microbiol, 2020, 22(7): 2792–2810.

[70] Wang X., Zha W., Liang L., Fasoyin O.E., Wu L., Wang S. The bZIP Transcription Factor AflRsmA Regulates Aflatoxin B(1) Biosynthesis, Oxidative Stress Response and Sclerotium Formation in Aspergillus flavus [J]. Toxins (Basel), 2020, 12(4).

[71] Vig I., Benkő Z., Gila B.C., Palczert Z., Jakab Á., Nagy F., Miskei M., Lee M.K., Yu J.H., Pócsi I., Emri T. Functional characterization of genes encoding cadmium pumping P(1B)-type ATPases in Aspergillus fumigatus and Aspergillus nidulans [J]. Microbiol Spectr, 2023, 11(5): e0028323.

[72] Zaccaria M., Ludovici M., Sanzani S.M., Ippolito A., Cigliano R.A., Sanseverino W., Scarpari M., Scala V., Fanelli C., Reverberi M. Menadione-Induced Oxidative Stress Re-Shapes the Oxylipin Profile of Aspergillus flavus and Its Lifestyle [J]. Toxins (Basel), 2015, 7(10): 4315–4329.

[73] Criddle D.N., Gillies S., Baumgartner-Wilson H.K., Jaffar M., Chinje E.C., Passmore S., Chvanov M., Barrow S., Gerasimenko O.V., Tepikin A.V., Sutton R., Petersen O.H. Menadione-induced reactive oxygen species generation via redox cycling promotes apoptosis of murine pancreatic acinar cells [J]. J Biol Chem, 2006, 281(52): 40485–40492.

[74] Ferreira G.F., Baltazar Lde M., Santos J.R., Monteiro A.S., Fraga L.A., Resende-Stoianoff M.A., Santos D.A. The role of oxidative and nitrosative bursts caused by azoles and amphotericin B against the fungal pathogen Cryptococcus gattii [J]. J Antimicrob Chemother, 2013, 68(8): 1801–1811.

[75] Caceres I., Khoury A.A., Khoury R.E., Lorber S., Oswald I.P., Khoury A.E., Atoui A., Puel O., Bailly J.D. Aflatoxin Biosynthesis and Genetic Regulation: A Review [J]. Toxins (Basel), 2020, 12(3).

[76] Sun Q., Shang B., Wang L., Lu Z., Liu Y. Cinnamaldehyde inhibits fungal growth and aflatoxin B1 biosynthesis by modulating the oxidative stress response of Aspergillus flavus [J]. Appl Microbiol Biotechnol, 2016, 100(3): 1355–1364.

[77] Caceres I., El Khoury R., Bailly S., Oswald I.P., Puel O., Bailly J.D. Piperine inhibits aflatoxin B1 production in Aspergillus flavus by modulating fungal oxidative stress response [J]. Fungal Genet Biol, 2017, 107: 77–85.

[78] He Z.M., Price M.S., Obrian G.R., Georgianna D.R., Payne G.A. Improved protocols for functional analysis in the pathogenic fungus Aspergillus flavus [J]. BMC Microbiol, 2007, 7: 104.

[79] Hong S.Y., Roze L.V., Wee J., Linz J.E. Evidence that a transcription factor regulatory network coordinates oxidative stress response and secondary metabolism in aspergilli [J]. Microbiologyopen, 2013, 2(1): 144–160.

[80] Joshi A.S., Cohen S. Lipid Droplet and Peroxisome Biogenesis: Do They Go Hand-in-Hand? [J]. Frontiers in Cell and Developmental Biology, 2019, 7.

